# Molecular dynamics simulations of intrinsically disordered protein regions enable biophysical interpretation of variant effect predictors

**DOI:** 10.1101/2025.05.07.652723

**Authors:** Aziz Zafar, Chao Hou, Naufa Amirani, Yufeng Shen

## Abstract

Predictive models for missense variant pathogenicity offer little functional interpretation for intrinsically disordered regions, since they rely on conservation and coevolution across homologous sequences. To understand the extent to which biophysics modulates model performance compared to genomic conservation, we model biophysics of IDRs explicitly for improved interpretation of variant effects. We develop MDmis, a method that uses biophysical features extracted from molecular dynamics (MD) simulations of IDRs to predict pathogenicity. We find that pathogenic variants in Long IDRs manifest differently, with transient order and depleted solvent access, compared to those in Short IDRs. Using MD simulations of sequences with single missense variants, we identify stronger evidence for pathogenic effects in Long IDRs compared to Short IDRs. MDmis, when combined with conservation information, achieves strong predictive accuracy of pathogenicity of variants in Long IDRs. Overall, extracting information from MD simulations can help understand the drivers of predictive performance and elucidate biophysical behaviors affected by pathogenic variants.

## Introduction

Accurate classification of missense variant effect is critically important in analysis of rare variants in genetic studies and clinical testing. More so, for developing therapeutic interventions and achieving precision genomic medicine, it is imperative that such variant effect predictions are biologically interpretable. However, a vast majority of variants are labeled of “unknown significance”, and even fewer are investigated for functional effect. Models used to predict missense variant pathogenicity can be pivotal to our understanding of protein function as it pertains to structured domains and disordered regions.

To date, there are many accurate models that can accurately predict proteome-wide missense variant effect^1–6^. These models primarily use conservation and co-evolution in multiple-sequence alignments (MSAs), with some structural and physiochemical features of amino acids and proteins, to make accurate predictions. Since pathogenic variants are under greater selection pressure, they are nearly absent in MSAs of homologous sequences. Therefore, by using information derived from MSAs, these models rely on a consequence of pathogenicity to predict pathogenicity itself, instead of modeling a purely biophysical cause of the mutation’s effect. As a result, there is a growing need to use explicit models of protein biophysics to jointly investigate cause, effect, and consequence of missense variants. Furthermore, since ordered and conserved domains are well-represented in homologous sequences, these models may excel at ordered regions but systematically underperform for a subset of intrinsically disordered regions (IDRs) and have stark incongruence^7–9^. This happens for two key reasons: 1) IDRs do not fold into stable structures and are highly dynamic and 2) IDRs evolve rapidly in their primary sequences across protein families^10–12^.

Although less structured and less conserved than ordered regions, IDRs play many key biological roles, such as cell signaling, protein regulation, complex formation, and RNA binding^13–15^. IDRs perform their cellular functions by being biophysically conserved, such that ensemble properties of IDRs tend to be under selection. IDRs typically exist as tails, short loops, longer linkers, sometimes very long (800-1400 residues) exterior shells, and even entirely disordered proteins (IDPs, which we will refer to as IDRs).

IDRs tend to have conserved properties such as their end-to-end distance, radius of gyration, compaction, and importantly, their abilities to phase separate and undergo disorder-to-order transition^12,16–18^. Recent studies have done much work to outline that IDRs are conserved in sequence space differently than ordered regions in three key ways: 1) IDRs have conserved amino acids composition, so missense variants may occur but will favor residues with similar physiochemistry 2) IDRs have conserved charge patterning, since increased segregation of charge results in compact proteins and uniform charge distribution causes less compact proteins, and 3) Some IDRs have motifs for post-translational modifications, specifically phosphorylation, since phosphate groups can modulate structural and functional changes to the protein^17,19–23^. Compositional bias has been found to be correlated with phase-separating and nuclear binding IDRs^24^. End-to-end distance and compaction are likely modulated by charge patterning and are especially important for linker IDRs to ensure that two ordered domains can be brought closer or kept apart depending on the global function of the protein^17^. In addition to ensemble properties of IDRs, such as compaction and transient order, IDR-mediated phase separation is key to protein functions, such as cellular fitness and transcription^25,26^.

Some work has been done to incorporate IDR-specific features, such as phase separation and ensemble properties, for more accurate prediction of missense variant effect and other fitness landscape tasks^8,27,28^. However, using only phase separation and changes in radius of gyration is limiting since these properties are affected in a subset of pathogenic missense variants. Instead, there is a need for inverting the study and using labels of pathogenicity to uncover properties of IDRs implicated in pathogenic variants. MD simulations can capture interaction dynamics, ensemble properties, and phase behavior of IDRs^28–31^. Compared to experimental ensembles generated using Nuclear Magnetic Resonance (NMR) Spectroscopy, MD simulations allow smaller timesteps down to 2 femtoseconds and capture fine scale dynamics such as side-chain rotations and transient structure^32,33^. Therefore, we decided to leverage interpretable features derived from molecular dynamics (MD) simulations to predict pathogenicity of missense variants. Our goal is to understand biophysical underpinnings of pathogenic variants in IDRs by comparing AlphaMissense, ESM1b, and a model trained on interpretable features derived from molecular dynamics simulations, named MDmis. MDmis is a workflow that uses a comprehensive set of features derived from MD simulations, such as surface area solvent access (SASA), residue fluctuation, secondary structure, angles, bonding interactions, and co-movement, to explicitly capture IDR ensembles and let trained machine learning models identify biophysical patterns of pathogenic variants. MDmis is trained on MD simulations of IDRs in the human proteome to predict benign and pathogenic labels from sources such as PrimateAI and ClinVar, among others^34–36^.

We found that MDmis can capture an IDR-wide signal of pathogenicity and provides marginal improvements when integrated with ESM1b for variant prediction in Long IDRs. We also find that pathogenic variants manifest differently based on IDR length. Long IDRs show disorder-to-order transition and depletion of surface area during their trajectories. We show that both behaviors are reversed by single pathogenic missense variants in MD simulations. On the other hand, Short IDRs have a weaker biophysical signal, and instead show strong enrichment of post-translational modifications, phase separation regions, and transcriptional-factor regulatory regions. Our results bring insight into the utility of MD simulations for understanding pathogenicity in IDRs and suggests distinct manifestations of pathogenic missense variants.

## Results

### Deriving features from MD simulations to train MDmis

MDmis uses coarse-grained molecular dynamics (MD) simulations of wild-type intrinsically disordered regions (IDRs). To extract features that can capture general biophysical characteristics of the IDR, we derive surface area solvent access (SASA), Root-Mean-Squared Fluctuation (RMSF), secondary structure assignments, chi, phi, and psi angles, residue movement covariance, and 9 bonding interactions. We also leveraged features extracted from mammalian MSAs, including charge pattern, compositional bias, and sequence entropy (Table **1**). Previous work has shown that these features can be used for rich embeddings of protein dynamics and can predict ensemble properties of IDRs such as radius of gyration^28^. Instead of using entire tensors, we isolate features of a window around the residue where mutations occur for supervised training of Random Forests (RFs). Using these representations and model predictions, we investigate potentially functional effects of pathogenic variants and follow-up using MD simulations of IDR sequences with single missense variants (see Methods and Figure **1**).

**Figure 1.**
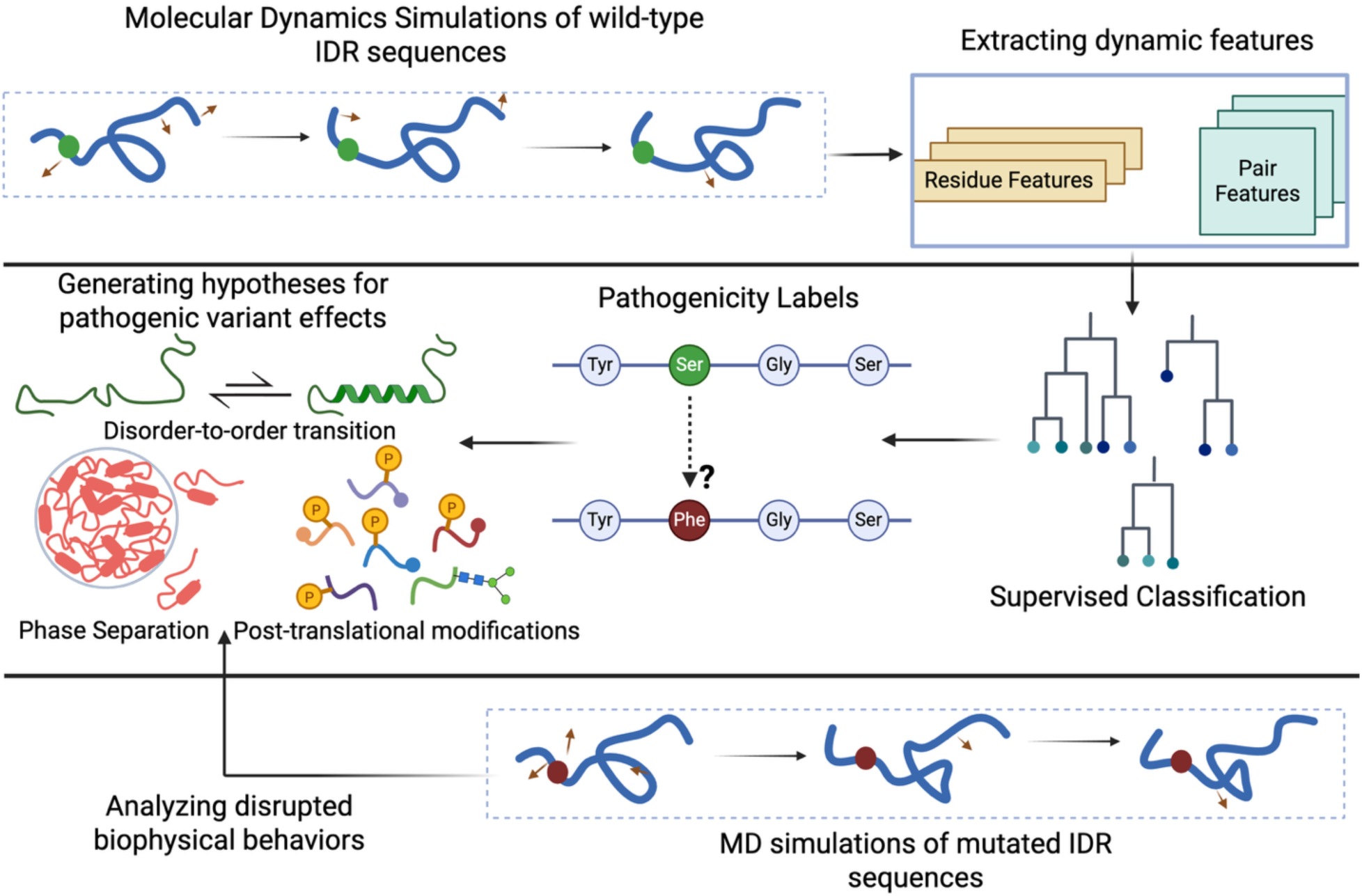
Workflow of data extraction, training, and analysis for MDmis. MDmis is a comprehensive approach for extracting explicit and interpretable dynamic features from molecular dynamics simulations of wild-type sequences. These features can be used for accurate prediction of pathogenicity in IDRs and for investigating potential functional effects of pathogenic variants.

**Table 1.** Features used in MDmis training regime. Residue features capture residue level attributes of mutated site (window size of 1) and leveraged 3 on both sides of the mutation (window size of 7). Pair features capture the interactions of the mutated residue with the rest of the IDR. Multiple-sequence alignment (MSA) features are extracted from Zoonomia’s mammalian genomes, with an emphasis on entropy, sequence compositional bias, and local charge pattern considering 9 amino acids on either side of the mutated residue. *Bonding interactions were only considered if a pair of residues had a certain bond for greater than 40% of the trajectory.

| Feature Group | Feature Name | Feature Value | Window Size |
| --- | --- | --- | --- |
| Residue | Solvent Accessible Surface Area | Mean | 1 or 7 |
|  | Solvent Accessible Surface Area | Standard Deviation | 1 or 7 |
|  | Root-mean-square fluctuation | Value Normalized by Mean | 1 or 7 |
|  | 8 Secondary Structure Assignments | Proportion of frames | 1 or 7 |
|  | 12 Percentiles of Chi Angles | Proportion of frames | 1 or 7 |
|  | 12 Percentiles of Phi Angles | Proportion of frames | 1 or 7 |
|  | 12 Percentiles of Psi Angles | Proportion of frames | 1 or 7 |
| **Pair** | 9 Bonding Types | Number of Interactions* | 1 |
|  | Covariance | Mean z-scored value | 1 |
| **MSA** | Sequence Composition | Average Cosine Similarity | 18 |
|  | Region Entropy | Average | 18 |
| | Charge Pattern ($\gamma$) | Average | 18 |
| | Charge Pattern ($\gamma$) | Standard Deviation | 18 |

### Benchmarking models in IDRs reveals predictive bias of conservation-based models

We first investigate if MDmis can capture a biophysical signal of pathogenicity by deriving information from wild-type MD simulations. We split 8,531 proteins with 12,303 IDRs at the protein level using 5-fold cross validation. For our evaluation metric, we use AUROC-0.1, which considers the receiver operating characteristic (ROC) curve only for false positive rate under 0.1, thereby shifting the baseline of random chance predictions to 0.05 (see Methods). Our trained models achieved an average AUROC-0.1 score of 0.29, revealing that explicitly modeling pathogenicity as a function of protein dynamics or even molecular effect of variant effect can yield reasonable performance in IDRs in the absence conservation information. AlphaMissense and ESM1b continued to be stronger performing models, indicating that conservation information continues to outperform explicit biophysical modeling of single residues (Figure **2A**). This result raised the possibility that genomic conservation may be strongly associated with pathogenicity in IDRs. We used GERP RS++ scores to measure genomic conservation across species, with a score >2 indicating a highly conserved site. We found a clear pattern demonstrating that GERP RS++ score was higher for sites with pathogenic variants in IDRs and in other protein regions (p<0.0001), with IDRs exhibiting a qualitatively clearer separation (Figure **2B**). Upon further analysis of predictive models on highly conserved sites (GERP RS++ > 2) and poorly conserved sites (GERP RS++ <= 2), we found that all models underperformed on poorly conserved IDR sites (Figure **2C**, **2D**). Previous studies have showed that conservation-based models including AlphaMissense and ESM1b can reach AUROC scores beyond 0.9 on certain datasets, but we see lower performance on variants in IDRs^2,37^. We confirm that this is likely due to widespread patterns of lower genomic conservation of sites in IDRs compared to other protein regions (Supplementary Figure 1A). Focusing on AlphaMissense scores, we confirmed that its predicted output is likely driving its performance in IDRs, such that only sites with very high GERP RS++ scores receive high AlphaMissense probabilities close to 1. However, in primarily structured protein regions, a great number of highly conserved sites were labeled by AlphaMissense as likely benign, likely implying that structural context is more relevant in non-IDRs for the predictions (Supplementary Figure 1B). In IDRs, AlphaMissense’s main source of information is likely conservation, due to a lack of structural context.

**Figure 2.**
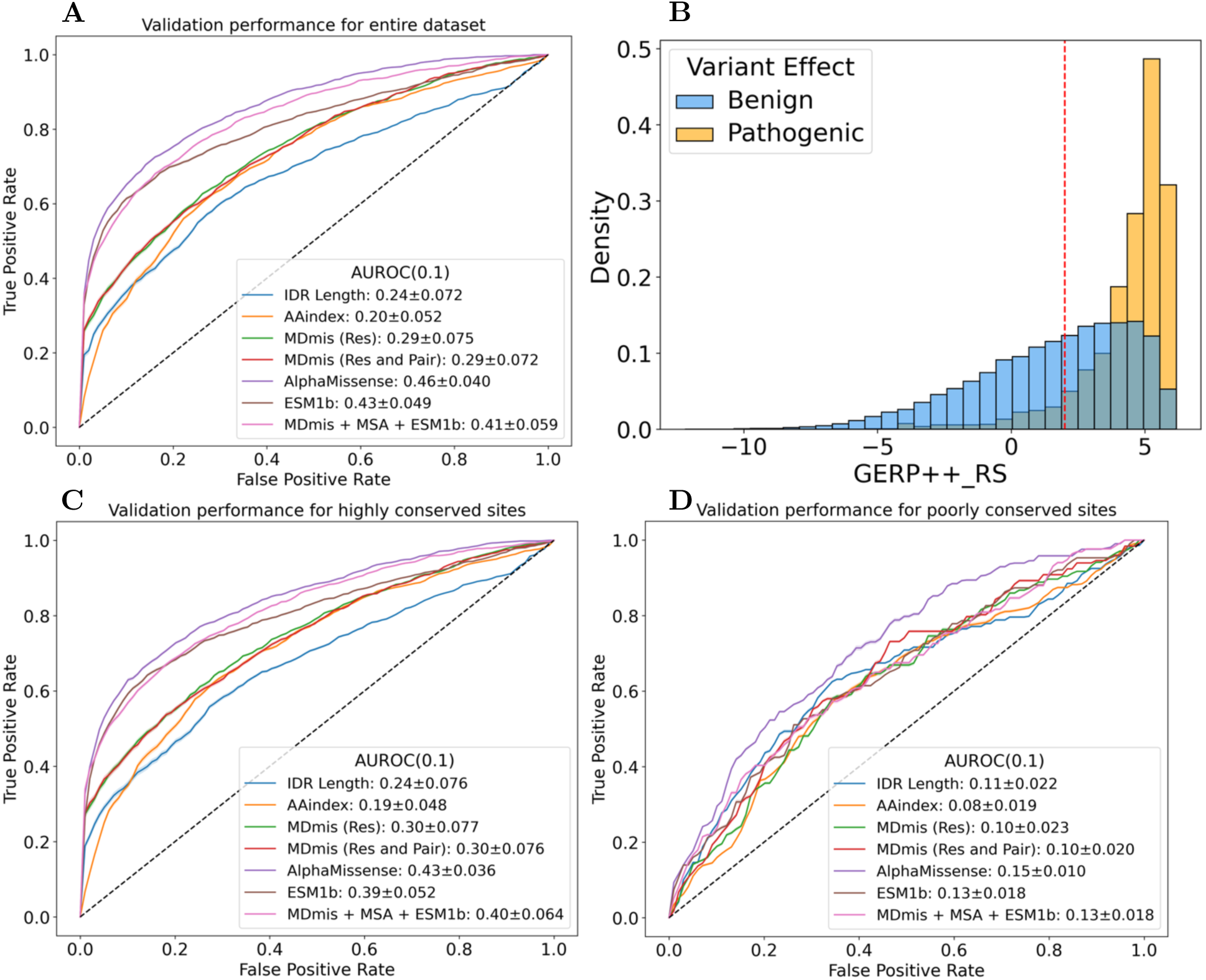
Genomic conservation of pathogenic training labels bias modulate performance in IDRs. (**A)** We compare the performance of MDmis, using various feature sets, with AlphaMissense and ESM1b on 13,340 Benign and 1,728 Pathogenic Variants. **(B)** In IDRs, pathogenic variants show a transposed distribution of GERP++ RS scores, which measures conservation across mammalian reference genomes. **(C)** Probing how models perform variants in highly constrained sites, with GERP++ RS>2, most models see improve performance. For this evaluation, 7,219 Benign and 1,586 Pathogenic variants were used. **(D)** On the other hand, on poorly conserved sites, all models demonstrate much poorer performance. For this evaluation, we used 6,121 Benign and 142 Pathogenic variants. All AUROC scores are computed for False Positive Rate<0.1 and multiplied by 10. AUROC-0.1 for random chance is 0.05. Scores are shown as mean and standard error of the mean for 5 testing folds.

To investigate the role of order and pathogenicity in IDRs, we compared pLDDT scores from AlphaFold2 across pathogenic and benign variants. Pathogenic variants showed a previously studied pattern of having higher order, measured using pLDDT (Supplementary Figure 1C). While this difference was also significant in other protein regions, we attributed this difference due to inherent variation in pLDDT present in the other protein regions. We further restricted our attention to MD simulations from G-protein coupled receptors (GPCRs), which are typically ordered. GPCRs were found to have a peak of pLDDT at 90 for all variants and our MD simulations showed that residues in GPCRs spent an average of 60% of their trajectory as alpha-helices, qualifying them as a positive control (Supplementary Figures 3C). We found that in GPCRs, pLDDT was not significantly different between pathogenic and benign variants (Supplementary Figure 3B). These findings indicate that levels of disorder likely play a vital role in pathogenic variants in IDRs.

### Comparison of models shows disparities across deep mutational scanning assays

We compared how AlphaMissense, ESM1b, and MDmis generalize to unbiased to 1,697 labels obtained from deep mutational scans of 24 proteins. Since typical DMS assays focus on ordered domains due to higher throughput, we were left with few labels and sites in total. Additionally, most of variants are in the peripheries of ordered regions, such that we had to use MDmis with single residue features instead of the window of 7 around a mutant site. While MDmis failed to generalize to deep mutational scanning data, with statistically insignificant Spearman rho correlations, it showed reasonable performance for activity and binding assays. As for the other models, we saw a strong decrease and disagreement in performance of conservation-based models on IDRs across assay types, compared to entire assays. ESM1b scores had strongest correlation with damage readouts for activity assays, whereas AlphaMissense scores had little to no correlation. On the other hand, both ESM1b and AlphaMissense have strong performance of assays measuring protein abundance. For binding assays, conservation-based models exhibit inverse correlation (Table **2****A**). However, when performances of ESM1b and AlphaMissense were considered on the entire assays, except for starting methionine residues, they showed mostly positive correlations, as expected based on AlphaMissense’s benchmark on ProteinGym assays (Table **2****B**). For binding assays specifically, both models showed lower correlation, with ESM1b still having negative correlation (*ρ* = -0.089, p<0.0001). Overall, this shows that due to the biases in training sets towards ordered and conserved protein regions, models are likely failing to generalize to unbiased held-out samples in IDRs. Additionally, binding assays could be particularly difficult for conservation-based models to capture.

**Table 2.** Hold-out testing of models indicate decreased generalization in IDRs. **(A)** Experimental readouts from deep mutational scanning (DMS) assays and their assay types are obtained from ProteinGym. DMS performance was evaluated only on residues and their single mutations that overlap with IDRome. All p-values for Spearman rank-correlation tests are provided in parentheses. **(B)** For this positive control, DMS performance was evaluated on single mutations for the entire screening assay, except for starting methionine residues. Spearman Rho correlation is shown between probability scores of models and experimental readouts multiplied by -1 to indicate molecular damage. Expression assays measure protein abundance. All p-values for Spearman rank-correlation tests were significant (p<0.0001).

| Assay Type | N | AlphaMissense | ESM1b | MDmis (Single Residue + Pair) |
| --- | --- | --- | --- | --- |
| Organismal Fitness | 935 | 0.11 (0.001) | -0.06 (0.52) | -0.014 (0.66) |
| Activity | 425 | -0.0098 (0.84) | 0.099 (0.041) | 0.083 (0.087) |
| Expression | 247 | 0.33 (<0.0001) | 0.37 (<0.0001) | 0.053 (0.40) |
| Binding | 90 | -0.28 (0.0084) | -0.58 (<0.0001) | 0.12 (0.28) |

**Table 2. Hold-out testing of models indicate decreased generalization in IDRs. (A)**
| Assay Type | N | AlphaMissense | ESM1b |
| --- | --- | --- | --- |
| Organismal Fitness | 80,882 | 0.18 | 0.22 |
| Activity | 44,886 | 0.36 | 0.33 |
| Expression | 14,182 | 0.48 | 0.49 |
| Binding | 4,123 | 0.13 | -0.089 |

### Features from molecular dynamics reveals functional length dichotomy of pathogenic variants in IDRs

We investigated the features that were used in training MDmis, especially those with greater feature importance. Using a baseline random forest that uses IDR length as a singular input, we observed that there was some predictive signal (Figure **2A**). Upon investigating the distribution of IDR length, we saw a right skewed distribution with very few IDRs beyond 500 amino acids (Supplementary Figure 2A). Given that pLDDT, a proxy of order was found to be associated with pathogenicity, we considered if we would find a similar result root-mean-squared fluctuation (RMSF), a MD based measure of disorder. However, RMSF for very long proteins can be misrepresentative, since longer proteins have greater frame-to-frame deviation simply by aligning backbones. We confirmed a clear linear relationship between length and unnormalized RMSF, indicating that due to the large spread of IDR length, RMSF is not reflecting fluctuation but rather a technical consequence of length (Supplementary Figure 2B). This is further supported by comparing pLDDT with RMSF in GPCRs, observing an expected inverse correlation (Supplementary Figure 3B). Compared to the IDRome’s wide distribution of lengths, the length of protein regions simulated in GPCRmd was on average 313 amino acids with a maximum length of only 495 amino acids.

Since region length is strongly correlated with residue fluctuation, we checked that bimodality in residue fluctuation was correlated with IDR length (Figure **3A**). Furthermore, as length of IDRs was used to determine the simulation time in IDRome, we wanted to account for the role of simulation time as a confounding variable. We selected ten Long IDRs for much shorter simulations of 150ns and confirmed that the average RMSF of the shorter simulations had a mean decrease of 0.613 (t = -7.11, p < 0.0001). However, since RMSF ranges for IDRs are varying from 0.5 Angstroms to 8 Angstroms, a decrease of 0.613 cannot solely explain the large difference in the observed two peaks.

**Figure 3.**
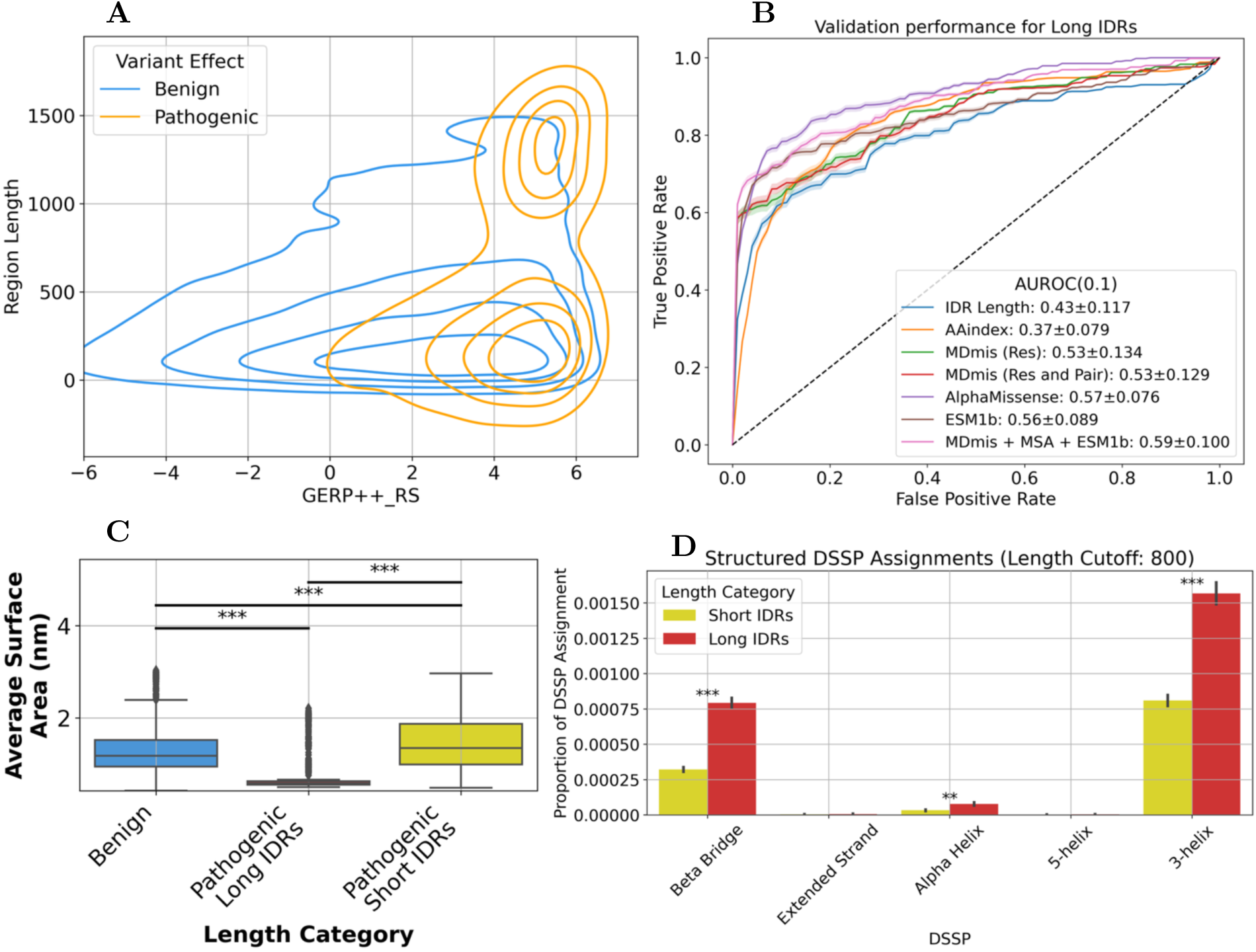
Pathogenic variants in Long IDRs are predicted with strong accuracy and have dynamic functions. **(A)** Density plots showing GERP++ RS score as a measure of genomic conservation and IDR length on the y-axis, colored by variant effect labels. Using an empirical cutoff of 800 amino acids to separate the two modes of pathogenic variants, we classify pathogenic variants in IDRs into Long versus Short. This yields 1,256 Pathogenic Short IDR variants and 636 Pathogenic Long IDR variants, with 13,340 Benign variants used for testing. **(B)** All predictive models show stronger performance, with MDmis, MSA, and ESM1b combined showing highest recall. Strong performance of biophysical models indicate that these variants likely impact a dynamic property of the residue. **(C)** Solvent accessible surface area (SASA) shows clear depletion for variants in Long IDRs. Short IDRs, on the other hand, have increased average SASA compared to benign variants. **(D)** Long IDRs also have increase proportion of ordered secondary structure assignments from DSSP. Beta Bridges, 3-helix, and alpha-helix are statistically significant with Mann Whitney U tests. All AUROC scores are for FPR<0.1. Scores are shown as mean and standard error of the mean for 5 testing folds. Error bars indicate 68% confidence interval (1 SEM). Significance stars: p<0.0001: ***, p<0.001: **, p<0.05: *

When variants are considered in tandem with the distribution of IDR length, we observe an underlying bimodal shape. We speculate that a few proteins in the right tail are enriched for pathogenic variants. Some of these “Long” IDRs are part of collagen proteins such as COL4A3, COL4A4, COL4A5, COL2A1, COL3A1, and COL4A1 and have an over-representation of pathogenic variants in the variant databases we used, including ClinVar and cancer hotspots. A total of 62 Long IDRs form this set, with 9 of them having more than 10 pathogenic variants each (Table **3**). Proteins with higher Mis-z scores and smaller o/e ratios have greater number of pathogenic variants and have lengths between 1300 to 2300^38^.

**Table 3.** Top 9 Long IDRs and their characteristics. IDRs were classified as Long empirically, based on the two modes appearing in the length density, as length >=800 amino acids. Missense z scores and o/e scores are taken from gnoMAD v4.1.0, except COL4A5 for which gnomAD v2.1.1 was needed. o/e measures the ratio of observed vs expected missense variants, and z-score acts as its significance level. Mis-z is greater for proteins with greater number of pathogenic variants in their Long IDRs and is insignificant for a few Long IDRs with fewer pathogenic variants.

| Protein | Gene | Region | Length | # Pathogenic Variants | Mis-z (o/e) |
| --- | --- | --- | --- | --- | --- |
| P29400 | COL4A5 | 1–1456 | 1685 | 131 | 2.5 (0.72) |
| P02458 | COL2A1 | 78–1254 | 1487 | 110 | 5.39 (0.67) |
| P02462 | COL4A1 | 1–1436 | 1669 | 61 | 4.99 (0.71) |
| P08123 | COL1A2 | 1–1134 | 1366 | 60 | 3.55 (0.77) |
| P02461 | COL3A1 | 79–1230 | 1466 | 47 | 4.61 (0.71) |
| Q01955 | COL4A3 | 1–1441 | 1670 | 35 | 2.46 (0.85) |
| P53420 | COL4A4 | 1–1456 | 1690 | 21 | 2.2 (0.87) |
| Q9UM47 | NOTCH3 | 1–1399 | 2321 | 21 | 1.87 (0.91) |
| Q5RHP9 | ERICH3 | 400–1530 | 1530 | 11 | –1.51 (1.1) |

### Features of MDmis implicate transient order and surface area depletion in Long IDRs

By empirically stratifying our IDRs into Long (>800 residues) and Short (<=800 residues), we probed how model performances change across these groups. Our Short IDR functional group has both very short (30-200 residues) and medium length (200-800) IDRs, but they are analyzed together since they appear as one mode in the length distribution of regions surrounding pathogenic variants. The distribution of variants for plotting is 1,256 Pathogenic Short IDR variants and 636 Pathogenic Long IDR variants, with 35,220 Benign variants. We first examined how AlphaMissense and ESM1b scores correlate in the context of these length groupings. We found that, albeit their scores were significantly correlated, the strength of correlation was not strong (*ρ*=0.45, p<0.0001). Additionally, ESM1b scores discerned pathogenic variants in Long IDRs more clearly compared to AlphaMissense. AlphaMissense, on the other hand, showed better separation between pathogenic variants in Short IDRs and Benign variants (Supplementary Figure 4).

Using ROC curves, MDmis alone had comparative to state-of-the-art performance on Long IDRs (Figure **3A**). Additionally, when MDmis was trained with Zoonomia MSA features and ESM1b scores integrated, the joint model outperformed other models with AUROC-0.1 of 0.59 (Figure **3B**).

This finding also suggested that pathogenic variants in Long IDRs have a greater signal from biophysical properties inferred from MD simulations. We probed the models’ top 10 features by Gini importance and found that solvent accessible surface area (SASA) ranked among the top 10 for both MDmis alone and with MSA and ESM1b integration (Supplementary Table **1)**. We further investigated SASA which was significantly decrease in Long IDRs, indicating that these residues are buried away from access to solvent (Figure **3C**). We also investigated the proportion of time each mutant residue spent in different ordered states and saw that residues in Long IDRs had a significantly higher chance of forming a 3-helix for 1 or 2 frames over the 1000 frames (Figure **3D**). We validated this using Hydrogen Bonds between backbones and found that residues in Long IDRs had significantly higher chances of having 3 bonds, indicative of a transient helical structure (Supplementary Figure 5B).

### Molecular dynamic simulations of mutated primary sequences support length dichotomy and mutational effects

Since MDmis was trained on MD simulations from wild-type protein sequences, it follows that biophysical properties that characterize pathogenic variants, in Long IDRs and potentially Short IDRs, may be disrupted by the variant. To test our hypothesis, we selected a random subset of variants from each group, and performed MD simulations with the mutated primary sequence using CALVADOS2^39^. We computed the same residue and pair-level features to identify a biophysical effect of the variant. For Long IDRs, we found that the ratio of frames as 3 Helix between the mutant and the wild-type simulation was significantly lower than Benign variants, indicating that the disorder-to-order transition is potentially disrupted by the variant (Figure **4A**). This was further supported by a peak difference in backbone-backbone hydrogen bonds of -3 after mutating residues in Long IDRs (Supplementary Figure 5B). Additionally, the solvent accessible surface area of the residue significantly increases with noticeable effect in long IDRs but not in Short IDRs (Figure **4B**). Consistently, in wild-type MD simulations, these residues were depleted for solvent access and potentially buried deep within the protein (Figure **3C**).

**Figure 4.**
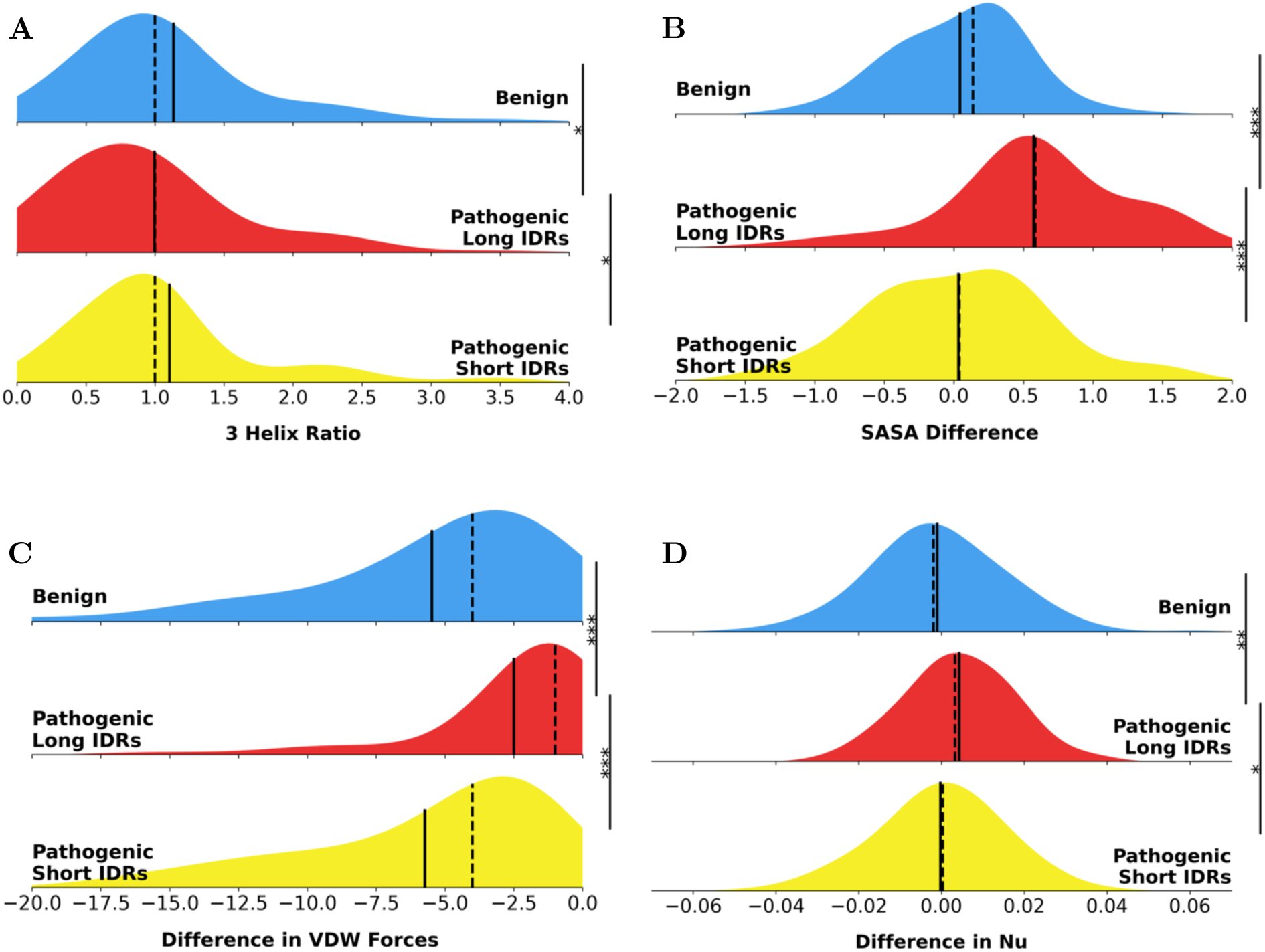
Comparing MD simulations of mutated IDRs shows disruption of dynamic behaviors in exclusively Long IDRs. By running MD simulations but changing single amino acids, we verified which dynamic features of wild-type IDRs are likely driving pathogenicity. **(A)** The ratio of 3-helix assignments between Long IDRs is lower compared to Benign variants and Short IDR pathogenic variants. **(B)** A similar pattern is observed, with much greater effect size and significance, for solvent access surface area (SASA), indicating that initial depletion of SASA is counter-acted by missense variants in Long IDRs. **(C)** These changes appear to manifest in Van Der Waal’s interactions for Long IDRs, but not for Short IDRs. **(D)** Additionally, compaction (nu) is also significantly increased in Long IDRs by single amino acid changes introduced by pathogenic variants. All analyses use Mann Whitney U Test. Significance stars: p<0.0001: ***, p<0.001: **, p<0.05: *. Mean: solid line, median: dashed line.

However, for Short IDRs, a mutation does not translate into statistically significant changes in biophysical properties, such as the number of VDW interactions or the compaction of the protein (Figures **4C****, 4D).** In contrast, the compaction of Long IDRs is changed by a single variant, indicating a much larger effect size on global ensemble properties via local residue level properties. These results further support that pathogenic variants in Long and Short IDRs manifest differently and have distinct functional effects.

### Sequence conservation and phosphorylation explain pathogenic variants in Short IDRs

Since Short IDRs do not have biophysical features affected by a mutation, we asked if Short IDRs possess a strong signal from wild-type MD simulations. We found that model performances in Short IDRs were generally lower and strongly favored conservation models such as AlphaMissense and ESM1b (Figure **5A**). We sought to investigate why sites in Short IDRs have a weaker signal from MDmis compared to sequence conservation from ESM1b. Our feature importances showed that MSA derived features, such as entropy of site and average entropy of window, were strongly predictive of pathogenic variants (Supplementary Table **1)**. We found that pathogenic variants in Short IDRs exhibit low median site-specific entropy comparable to Long IDRs, albeit the mean-rank is significantly greater for Short IDRs (Figure **5B**). However, there is a significant decrease in median entropy for the window around the mutation, compared to Long IDRs (Figure **5D**). We also observed a significant increase in charge segregation in the windows around a pathogenic variant in Short IDRs compared to their Long IDR pathogenic counterparts (Figure **5C**). Because charge segregation typically modules compaction, it followed that length adjusted compaction, nu, was also significantly higher for Short IDRs we compared to Long IDRs. In addition, the number of Van Der Waal’s (VDW) interactions during the trajectory were higher in Short IDRs. Together, these indicate a weaker explicit signal from biophysical features, but a stronger association with sequence properties that modulate ensemble behaviors.

**Figure 5.**
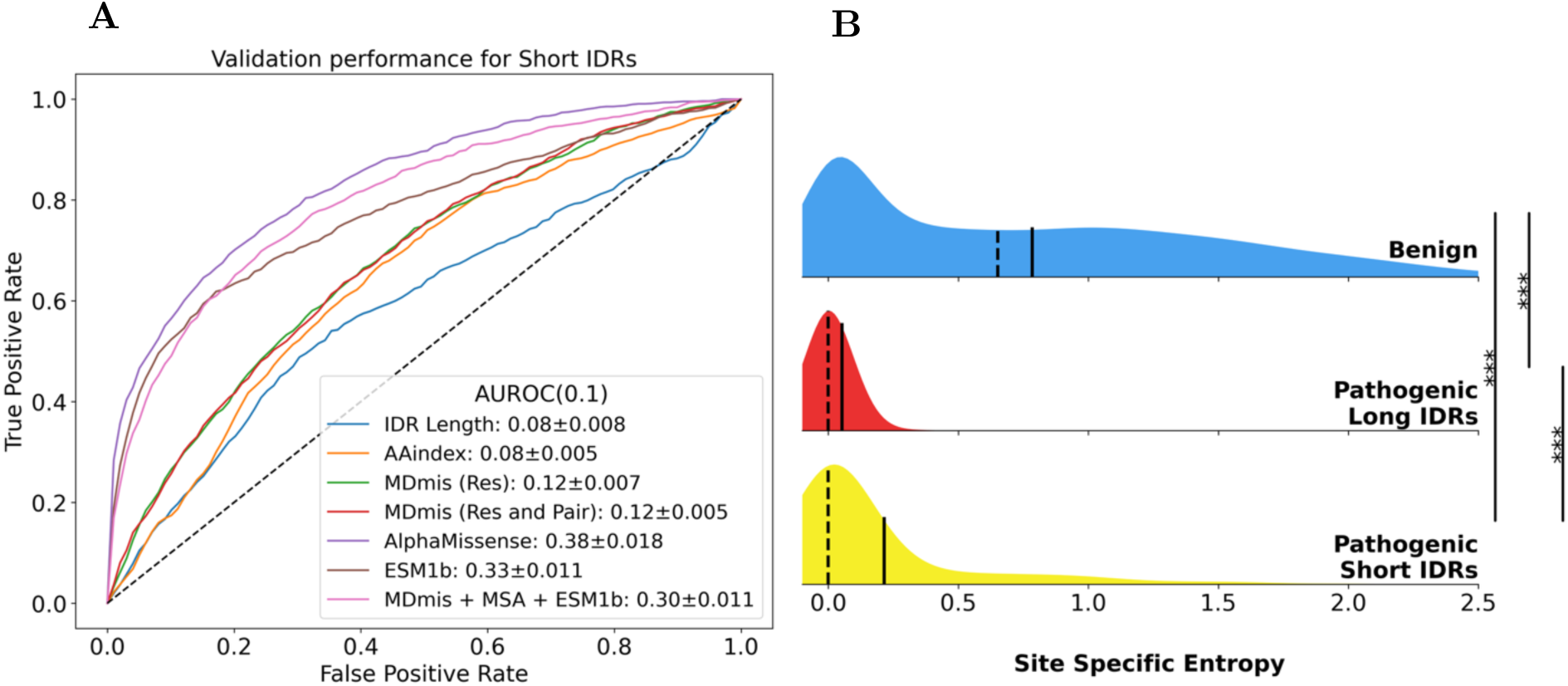
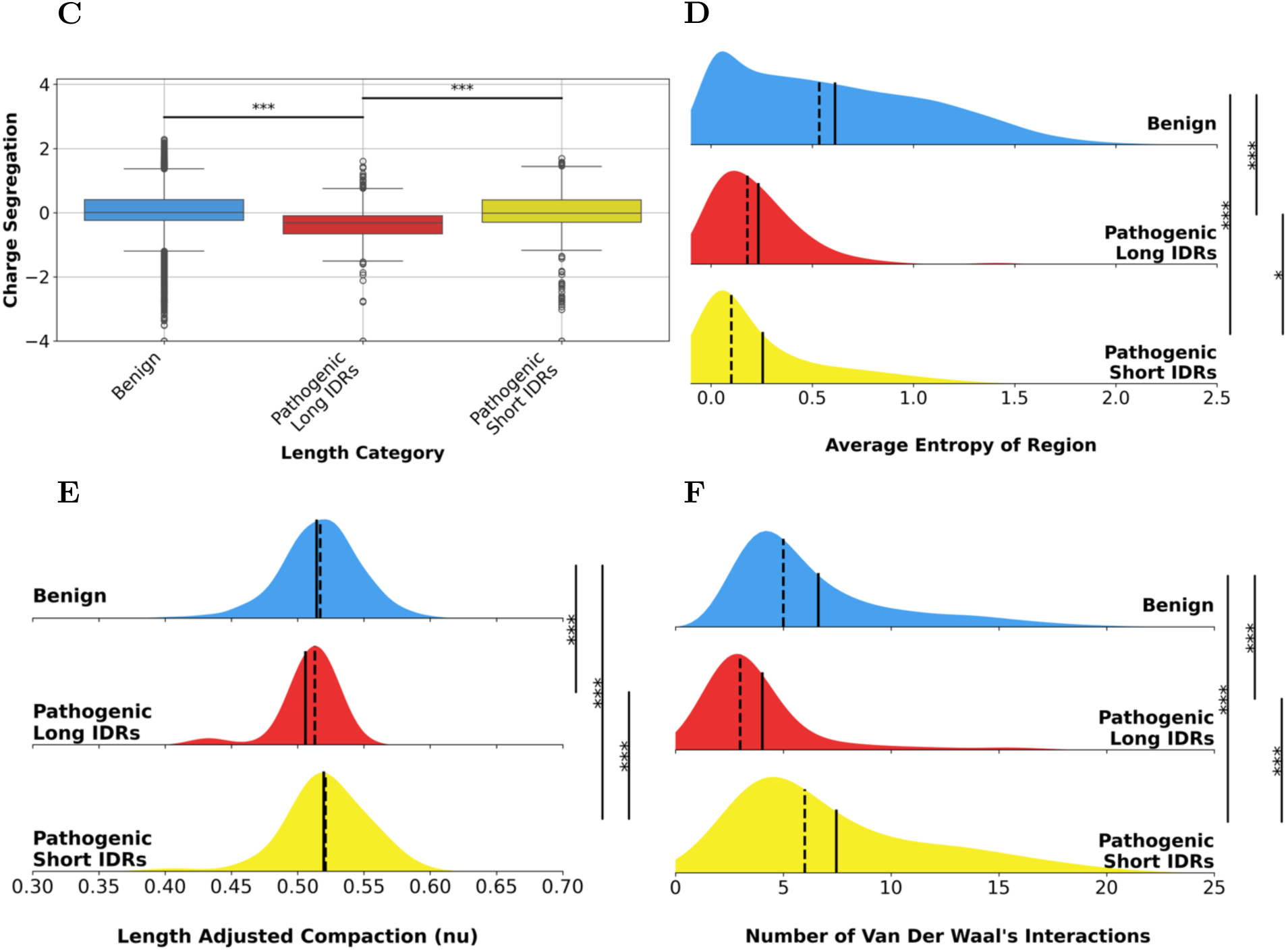
Pathogenic variants in Short IDRs are predicted better by evolutionary information and exhibit sequence-dependent dynamic properties. **(A)** When evaluating models on Short IDRs, MDmis shows weaker performance but can supplement ESM1b in regions of low precision. AlphaMissense and ESM1b show strongest performances, indicating that sequence properties of these variants are likely more important. **(B)** Short IDRs show lower site-specific entropy in Zoonomia MSAs compared to Benign variants. **(C)** Charge segregation is plotted in log-scale as log_10_(γ + epsilon), where γ measures segregation using small overlapping windows. Charge segregation is significantly higher in Short IDRs compared to Long IDRs. **(D)** Average entropies of short windows around variants show significant decrease for pathogenic variants in Short IDRs. **(E)** Length adjusted compaction measures overall compactness of the IDR using a scaling exponent. With a qualitatively small effect difference, pathogenic variants in Short IDRs are also more compact than Long IDRs and benign variants. **(F)** Number of Van Der Waal’s forces that occur in greater than 40% of the frames are significantly higher between variant site and other residues in Short IDRs. All analyses use Mann Whitney U Test. AUROC score is computed for FPR <0.1. Scores are shown as mean and standard error of the mean for 5 testing folds. Significance stars: p<0.0001: ***, p<0.001: **, p<0.05: *. Mean: solid line, median: dashed line.

Given that there is an absence of a transient secondary structure in Short IDRs, we questioned if this group is mostly composed of regulatory regions and modification motifs. It is well-studied that IDRs can be regulated by post-translational modifications, primarily via phosphorylation of Serine,

Threonine, and Tyrosine. We used dbPTM, a database of putative post-translational modifications (PTMs) to examine whether enrichment of PTMs was associated with Pathogenic variants in Short IDRs^40^ (Table **4****A**). We found that odds of identifying a PTM site in Short IDRs were significantly higher than in Long IDRs, with phosphorylation being the most common modification (OR =12.5, p <0.0001). We also investigated whether variants changing a Serine, Threonine or Tyrosine into other amino acids was enriched in Short IDRs and found an insignificant result (OR=2.8, p=0.29). This indicates that while phosphorylation motifs are likely enriched in Short IDRs, their sole disruption may not completely explain pathogenicity (Table **4****B**).

**Table 4.** Post-translational modifications are enriched in pathogenic variant sites in Short IDRs. **(A)** Curated post-translation modification sites were identified and overlapped with pathogenic variants. 89 PTMs that were identified in Pathogenic Short IDRs were phosphorylation sites. PTMs were overrepresented in Short IDRs. **(B)** STY: Serine, Threonine, and Tyrosine, are the three most commonly phosphyralted amino acids. STY to non-STY mutations, which could disrupt phosphorylated residues, were not significantly different between the two subtypes.

| **A** |  |  |
| --- | --- | --- |
| **Curated PTMs** | **Pathogenic**<br>**Long IDRs** | **Pathogenic**<br>**Short IDRs** |
| **None** | 631 | 1149 |
| **PTM site** | 5 | 114 |

**Table 4. Post-translational modifications are enriched in pathogenic variant sites in Short IDRs.**
| **B** |  |  |
| --- | --- | --- |
| **Amino Acid Change** | **Pathogenic – Long IDRs** | **Pathogenic – Short IDRs** |
| **STY to STY** | 3 | 18 |
| **STY to non-STY** | 10 | 168 |

Given that Van Der Waal’s force and Compaction are significantly different in Short IDRs, we questioned the role of phase separating regions, in addition to transcription-factor regulation domains. We observed that both phase separating regions in PhaSepDB and TF regulatory domains in TFRegDB are almost completely depleted in pathogenic variants in Long IDR^41,42^ (Table **5****A**). For regions that phase separate with the same proteins, there is a significant enrichment in Short IDR pathogenic variants (OR=3.04, p<0.0001). Similarly, regions that phase separate with other proteins are significantly overrepresented in Short IDRs (OR=1.86, p=0.016). As for TF regulatory domains, we observed that both activation and repression domains were also significantly enriched in Short IDRs (AD: OR =4.09, p<0.0001, RD: OR=2.37, p=0.00063), whereas bi-functional domains were found not to be represented in our variants (Table **5****B**). Altogether, these three functional annotations may explain pathogenic variants in Short IDRs. There is very little overlap in variant sites between these annotations, with at most 12 sites being shared between TFRegDB and PhaSepDB (Supplementary Figure 6).

**Table 5:** Regions involved in phase separation and transcription-factor regulation significantly correlated with pathogenic variants in Short IDRs. Curated regions involved in phase separation and regulation of transcription factors were identified and overlapped with all variants. **(A)** Phase separation with duplicates of the same protein as well as with other proteins were significantly overrepresented in Short IDRs, compared to Benign variants. Pathogenic variants in Long IDRs were completely missing phase separating regions. **(B)** Similarly, transcription factor domains, such as activation domains (AD) and repression domains (RD) are also significantly enriched in pathogenic variants in Short IDRs, with Long IDRs having clear depletion. Bi-functional domains (Bi-F) are not well-represented in many IDRs.

| Curated Phase Separation Regions | Benign | Pathogenic – Long IDRs | Pathogenic – Short IDRs |
| --- | --- | --- | --- |
| None | 34482 | 635 | 1190 |
| PS – Self | 457 | 0 | 48 |
| PS – Other | 281 | 1 | 18 |

**Table 5: Regions involved in phase separation and transcription-factor regulation significantly correlated with pathogenic variants in Short IDRs.**
| Curated TF Separation Regions | Benign | Pathogenic – Long IDRs | Pathogenic – Short IDRs |
| --- | --- | --- | --- |
| None | 34624 | 634 | 1186 |
| AD | 371 | 1 | 52 |
| RD | 222 | 1 | 18 |
| Bi-F | 3 | 0 | 0 |

## Discussion

The function and conservation of IDRs and their highly dynamic physical properties is still not well-understood. While missense variant predictors provide an opportunity to understand how certain residues play a role of IDR function, they might under-perform and offer little interpretation to the underlying biophysical effect to the protein. To address these issues, we performed an analysis using MD simulations of the human disordered proteome and its predictive signal for pathogenic missense variants. Our main goal was to compare an explicit model of IDR ensembles, MDmis, with conservation-based models to understand how biophysically damaging variants manifest in evolutionary models. We also aimed to use MD features to interpret the effect of variants on IDRs and then substantiated these hypothesized effects by generating ensembles of mutated IDRs.

We find that IDRs generally possess lower genomic conservation than ordered regions and that conservation scores in IDRs are very clearly predictive of pathogenicity. Additionally, we notice dramatic performance decreases in AlphaMissense, ESM1b, and MDmis in poorly conserved sites. However, without leveraging conservation information, we find that MDmis can capture a reasonably strong predictive signal and can improve upon ESM1b’s predictions.

In our analysis, we find strong evidence for a bimodal length effect in pathogenic variants, with Long IDRs having disorder-to-order transition and depletion of surface area. Conversely, Short (and medium length) IDRs have a much weaker biophysical signal, with significant differences mostly in sequence-dependent properties such as sequence entropy, charge segregation, and compaction. However, Short IDRs see a stronger correlation with phosphorylation motifs and regions involved in phase separation and transcription-factor regulation. Given the models’ much stronger performance on Long IDRs, we hypothesize that their transient structure and solubility are likely affected by pathogenic missense variants, similar to structured domains. By affecting residue level properties, these mutations may affect some global ensemble property of the IDR and the protein. We found that this subset of pathogenic variants is likely to adopt a 3-Helix structure during the trajectory. Additionally, residues are starkly depleted for solvent-access in Long IDRs, potentially due to the length itself. For longer IDRs, having some residues hide underneath a hydrophilic exterior shell is easier, compared to IDRs that are a few hundred amino acids and are likely linkers or tails. It is also possible that disorder-to-order transition into a helical structure prevents these residues from interacting with solvent, decreasing their average over the course of the trajectory. This may explain why mutating these residues affects both the 3-Helix proportion and SASA simultaneously. In doing so, the entire proteins compaction is also changed, possibly due to increased SASA and solvent mediated bonding interactions. These results indicate that Long IDRs are behaving analogous to ordered proteins for missense variant effect predictions.

Variants in Short IDRs, on the other hand, has a much weaker signal from MD simulations than sequence conservation. It is well-known that residues in IDRs can be conserved as phosphorylation sites for allosteric regulation^43,44^. We also find transcription-factor activation and repression domains over-represented in Short IDRs and absent in Long IDR pathogenic variants. Together, these regulatory functions can be vital in IDRs that belong to GPCRs and other receptor proteins. We also find strong evidence for Short IDRs overlapping with phase separating regions and transcriptionally regulating regions of the IDRs. However, since curated databases for PTMs, phase separation, and transcription-factor annotations are incomplete, it is difficult to speculate whether these functions are the sole explanations for pathogenic variants. Alternatively, we saw a significant difference in charge segregation and compactness, indicating that Short IDRs are less compact. Disordered regions tend to form short loops or medium length linker regions between structured protein domains. Therefore, we hypothesized that a variant affecting compactness, and the increased Van Der Waal’s interactions may be the biophysical cause of pathogenicity. However, in our in-silico mutated simulations, we did not see a significant change in either VDW forces or compaction. It is likely that these are properties important for the Short IDRs but are not disrupted by pathogenic variants. Overall, pathogenic variants in Short length IDRs remain a nebulous mixture and do not exhibit a unique and widespread signal of biophysical pathogenicity. For such variants, models that use homologous sequences and MSAs likely capture binding motifs and windows of residues under selection. Pathogenic variants in Long IDRs show a clearer pattern of disruption using MD simulations, from how accurately models can predict their effect to how mutated MD simulations reveal noticeable global biophysical changes from only a single amino acid change.

There are some key limitations in our findings that future work could address. Firstly, the average number of frames, while significantly greater for Long IDRs, was very low (1 or 2 per 1000). This may imply that these residues may become ordered transiently without any external factor but may not be stabilized unless there is DNA binding or complex formation^45,46^. For more rigorous understanding of stable disorder-to-order transition in these IDRs, running MD simulations with binding partners and co-factors would be vital^47,48^. Furthermore, while the IDRome database is a comprehensive catalog of IDR ensembles, it does not model IDRs with their ordered regions or with DNA, binding partners, ligands, or cofactors. IDRome and our simulated trajectories use coarse-grained MD simulations, which trade lower spatial resolution for longer simulations.

However, as All-Atom MD simulations become more tractable or CG force-fields start to incorporate ordered and disordered regions together, it will become easier to perform unbiased simulations of entire proteins with IDRs in their functional contexts. It is also likely that with more MD trajectories of proteins becoming accessible, our model could learn from MD simulations of both wild-type and mutated sequences for semi-supervised learning. Finally, clinical labels show bias towards regions of high genomic conservation and do not capture the range of functional effects. Therefore, training these models to learn patterns from allele frequencies may help uncover novel insights into IDR function and evolution^49^.

## Methods

### Input Molecular Dynamics trajectories

For the input to MDmis, we used the published IDRome database, consisting of MD simulations of 28,058 IDRs^34^. To avoid using conformationally unstable frames, we discarded the first 10 frames of each trajectory.

Coarse-grained simulations were converted to all-atom trajectories using cg2all^50^. To include MD simulations of ordered domains as a control, we used simulations of G-protein coupled receptors (GPCRs) from GPCRmd^51^.

We derived residue level features of shape L x 47 x 7, where L encodes the sequence length, and 47 features: mean and standard deviation of surface area solvent accessibility, root mean square fluctuation, proportion of 8 DSSP assignments, and proportion in 12 chi1, phi, and psi angle quantiles computed using mdTraj^52^. The residue features were considered for 3 sites to the left and right of the variant site. In case of variants at the edges, features were padded using all zeros. We derived features for each residue pair, with shape L x L x 10 such as covariance and 9 bonding interaction using GetContacts^53^. We also used the difference of 553 AAIndex features between original and changed amino acid for each variant, which is of shape 1 x 553 for each variant^54^.

### Zoonomia MSAs of IDRs

To extract IDR-specific conservation features, we leveraged the Zoonomia database, which has codon alignments of orthologous genes with hg38 as the reference. This database contains 447 animals, of which 241 are Zoonomia mammals and the remainder are primates^55^. Due to the evolutionary closeness to humans, we saw that IDRs were successfully aligned. We converted codon alignments to protein MSAs by converting triplets into corresponding amino acids and aligned the reference MSA to the matching UniProt sequence. We also removed lower quality alignments due to multiple orthologs, reducing depth to around 500 sequences per protein. We extracted average entropy of a window of 18 residues, 9 flanking on each side around every variant, excluding the variant site itself. We also computed entropy of the mutated site separately at the Site Entropy.

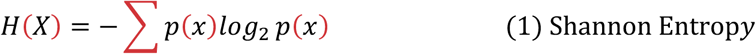

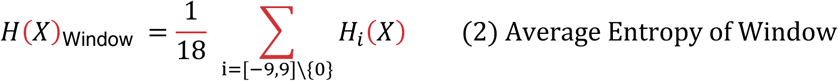

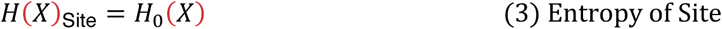

We computed conservation of sequence composition across the MSA by clustering amino acids using the first 5 principal components of their AAIndex features into 4 groups. Then, we count the number of residues in the window that are assigned to each of the 4 groups and compute the average cosine similarity of each sequence in the MSA with the UniProt reference, with a higher average indicating more conserved compositional bias.

Given a grouping of amino acids into four clusters and sequence j, for a window from -9 to 9 excluding 0

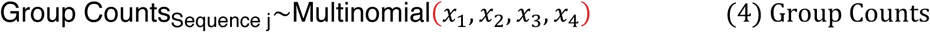

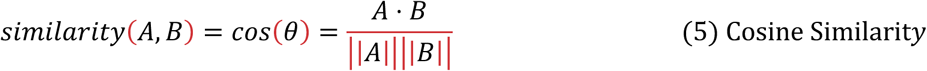

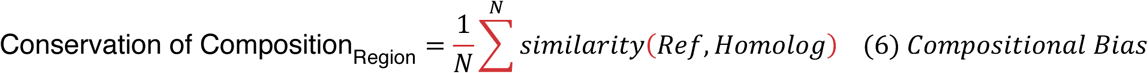

Here, N refers to number of homologs. Ref refers to the reference UniProt sequence and the homolog refers to the homologous sequences in the Zoonomia MSAs.

Lastly, we compute average and standard deviation of unscaled charge segregation γ, a precursor to the scaled metric *к* from Das and Pappu, 2013^20^. Higher gamma represents greater charge segregation, and a lower standard deviation indicates higher conservation of charge pattern in primates and mammals.

For 𝑁_segments_ overlapping segments of size *g,* we compute charge asymmetry for each *k* segment:

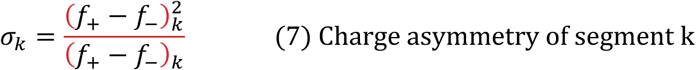

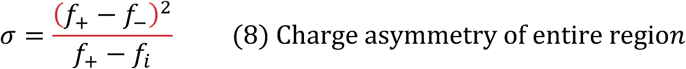

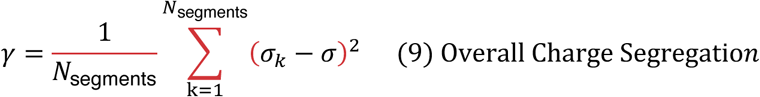

We do not scale the charge by the maximum segregation to obtain *к*, because use the conservation of charge pattern across homologous sequences as a feature. We also do not consider the entire IDR’s pattern because we want to leverage more precise resolution around the variant.

### Pathogenicity Labels

We used 37,118 curated and labeled missense variants from 12,488 unique protein regions overlapping with the IDRome database. 20,831 variants from PrimateAI and 14,392 from ClinVar were used as benign labels^35,36^. For pathogenic variants, 1074 variants from ClinVar, 405 variants from cancer hotspots, and 416 variants from other data sources were collected^46,56–58^. For ClinVar, all used labels were selected with at least one-star non-conflict submits and were filtered to remove start-loss mutations at the first methionine residue. However, upon integrating these data with features extracted from Zoonomia MSAs, 3 Pathogenic and 416 Benign variants were dropped as a result of missing MSA information. Our final training and data analysis was performed on 36,696 variants of 12,303 IDRs.

To include a background set of missense variants, we considered a larger set of variants obtained from ClinVar, PrimateAI, HGMD, cancer hotspots, and others, and removed all the variants used in training MDmis^59^. This set, labeled Other Protein Regions is not strictly but primarily ordered domains. It comprises 47,674 benign and 61,618 variants. We also subset variants in GPCRmd from this set, yielding 111 missense variants with 54 benign and 57 pathogenic variants of 41 unique GPCRs.

To perform hold-out testing, we used deep mutational scanning (DMS) assays taken from ProteinGym^60^. We multiplied the DMS readout with the provided directionality and -1 to convert readouts from fitness to damage. Assay types were provided in the ProteinGym metadata. We selected assays from the following 25 proteins manually to ensure at least 10 mutations that overlapped with IDRome: ADRB2 (P07550), CASP3 (P42574), CASP7 (P55210), CBS (P35520), CD19 (P15391), ERBB2 (P04626), GLPA (P02724), HMDH (P04035), KCNE1 (P15382), LYAM1 (P14151), MET (P08581), MSH2 (P43246), MTHR (P42898), PAI1 (P05121), PPARG (P37231), PPM1D (O15297), PRKN (O60260), PTEN (P60484), P53 (P04637), RAF1 (P04049), SC6A4 (P31645), SHOC2 (Q9UQ13), SYUA (P37840), TADBP (Q13148), YAP1 (P46937)^61–83^. By combining these data sources with our MD and MSA features, we retrieve 1,697 readouts. 935 of these readouts measure Organismal Fitness, 425 measure Activity, 247 measure Expression, and 90 measure Binding.

### Analysis of site-specific genomic conservation

To represent cross-species per-site conservation, we leveraged the GERP_RS++ score, taken from the dbNSFP database^84,85^. This score is taken from genome alignments of humans with other mammals to capture sites that are under high constraint i.e. with higher-than-expected rates of rejected substitutions. We aggregated the genome-level score into residue levels scores by taking an average of the scores of nucleotides in each codon. We defined a GERP_RS++ score > 2.0 as high constraint, and the converse as low constraint.

### Feature extraction and model training

We used a 5-fold cross validation based on unique proteins to avoid leakage between multiple IDRs of the same protein. We used combinations of the residue features at the site of mutation (titled Res), the average of covariance and the number of bonding interactions that exceed 40% of the trajectory (titled Pair), and the difference of 553 AAIndex features wild-type and mutant residue (titled AAIndex). 40% was chosen as an empirical cutoff to disregard potentially noisy interactions between residues. We also use sequence conservation features, such as charge pattern, standard deviation of charge pattern, sequence composition, and average entropy extracted from Zoonomia (titled MSA). Lastly, we passed ESM1b zero-shot LLR score as a feature our Random Forest models as an early integration ensemble model. Using these features, we train different Random Forest models using 100 random trees and maximum depth limited to 15 splits.

For training MDmis, pathogenic variants were split by protein ID. To ensure minimial leakage for AlphaMissense, benign variants from PrimateAI were used to train MDmis similar to AlphaMissense. Benign variants from ClinVar were used to evaluate performances for all predictors.

We compared the performance of each model with AlphaMissense and ESM1b^1,2^. To use ESM1b LLR score for receiver-operating characteristic curves, we used a min-max scaling of all LLR scores after multiplying them with -1. This allows greater LLR scores to be represented as close to 0 probabilities of pathogenicity. AlphaFold2 structures and pLDDT scores were used for analyzing disorder levels, in addition to qualitative structural analysis^86^.

### Model Evaluation

Area under receiver operating characteristic curve (AUROC) score typically describes the trade-off between true positive rate and false positive rate over all thresholds of the probability scores. We decided to compute the ROC values for each testing set and limited our analysis to thresholds where false positive rates are less than 0.1. Then, this score is multiplied by 10 to make it comparable to AUROC scores over all thresholds, although AUROC-0.1 has a random-chance performance of 0.05. We find that, in the absence of complete data where the true distribution of pathogenic and benign labels is known, AUROC-0.1 is likely a better estimate of model performance in settings where high precision is needed. AUROC-0.1 scores are shown as mean and standard error of the mean across 5 testing sets.

### Curated databases for protein type analysis

For post-translational modifications (PTMs), we used all PTM categories in the dbPTM database^40^. We did not remove sites that had more than one PTM, since dropping such duplicates did not change our analysis noticeably. For regions involved in phase separation, we used curated and annotated regions from PhaSepDB v2.1^41^. We did not use the few annotated entries where start and end loci of the regions were not clearly annotated, for instance “exon X” or “IDR of Protein Y”. Lastly, for transcription-factor regulation domains, we used TFRegDB, a set of curated human transcription-factor regulated domains^42^.

### Generating ensembles of mutated IDRs

We used CALVADOS2 to generate ensembles of mutated IDRs, using the same code and parameters used to generate IDRome^34,39^. C-alpha coarse-grained simulations were run using HOOMD-blue and OpenMM’s toolkit^87^. NVT ensembles were stabilized at 310 Kelvin with a Langevin Integrator with a time step of 10fs and friction coefficient of 0.01ps^’/^. Exact details for simulations and approach can be found in Tesei 2024^34^. IDR simulation time is scaled with length, with a minimum of 70ns and up to 6000ns. IDRs of length 30 – 40 that ran for 70ns required around 2 minutes to be simulated, whereas 1200 amino acids required around 6 hours on an RTX 4090 GPU. Due to long running times of Long IDRs and Benign variants, we selected random subsets of variants from each group for simulation in incremental batches. We used 229 simulations of benign variants, 196 variants of pathogenic variants in Long IDRs, and 562 simulations of pathogenic variants in Short IDRs. We decided to perform at least a 100 simulations per group to ensure representative samples per group and performed more simulations for Short IDRs due to their short runtime and biological interest.

To avoid dependence between frames and allow for stabilizing, 20 initial frames were discarded, and 800 random frames were selected from the remaining 980. Like our data from wild-type IDRs, coarse-grained simulations were converted to all-atom resolution before extracting the same 47 residue features at the exact site of mutation and 10 pairwise features used in MDmis^50^. We computed the difference between MD of mutated sequence and MD of wild-type sequence, as well as ratio for features measured as proportions, such as secondary structure assignments.

### Statistical analysis and plotting

Differences between medians were tested for statistical significance using Mann Whitney U Tests with Bonferroni correction in scipy^88^. For correlation analyses, Spearman’s Rho test was used, with visual assessment of monotonicity assumption. Chi-square statistics and corresponding p-values were computed with Yates correction. All plots were generated using seaborn and matplotlib^89,90^. Workflows and Venn diagrams were created and rendered in BioRender.

## Code and Data Availability

All scripts used in performing are available on our GitHub repository: https://github.com/ShenLab/MDmis.git. The tensors of features extracted from molecular dynamics features are available at https://huggingface.co/datasets/ChaoHou/protein_dynamic_properties for several MD databases, including IDRome, GPCRmd, and others. The processed data we used in our study and the coarse-grained simulations we performed are available on our Zenodo repository with DOI: 10.5281/zenodo.15346250.

## Author Contributions

Conceptualization: YS; Methodology: AZ, Software: AZ, CH; Formal analysis: AZ, NA; Data curation: AZ, CH; Writing – Original Draft: AZ; Writing – Review & Editing: YS, CH, NA; Visualization: AZ; Supervision: YS; Funding Acquisition: YS.

## Acknowledgements

This work is supported by grants from NIH (R35GM149527) and Simons Foundation Autism Research Initiative (SFARI #1019623).

## Declaration of interests

The authors declare no competing interests.

## Notes

https://zenodo.org/records/15346250

### Competing Interest Statement

The authors have declared no competing interest.

