## Supplemental Figures for "Molecular dynamics simulations of intrinsically disordered protein regions enable biophysical interpretation of variant effect predictors"

**A**

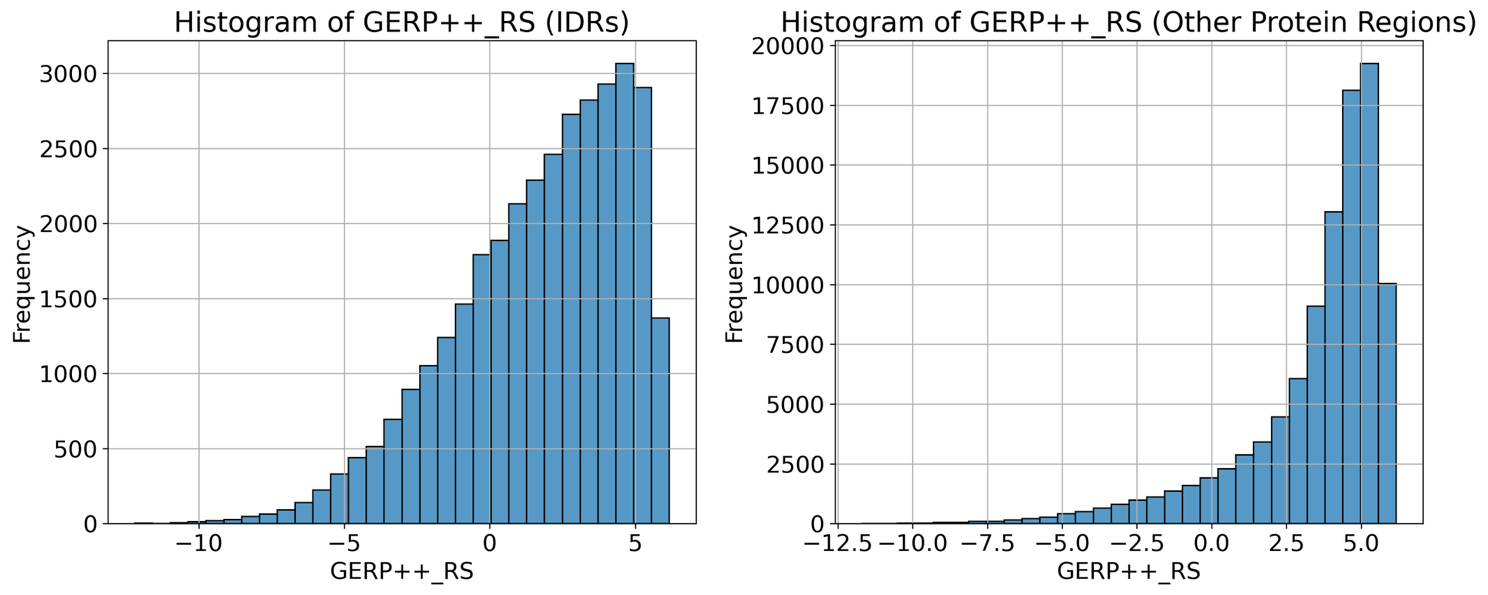

**B**

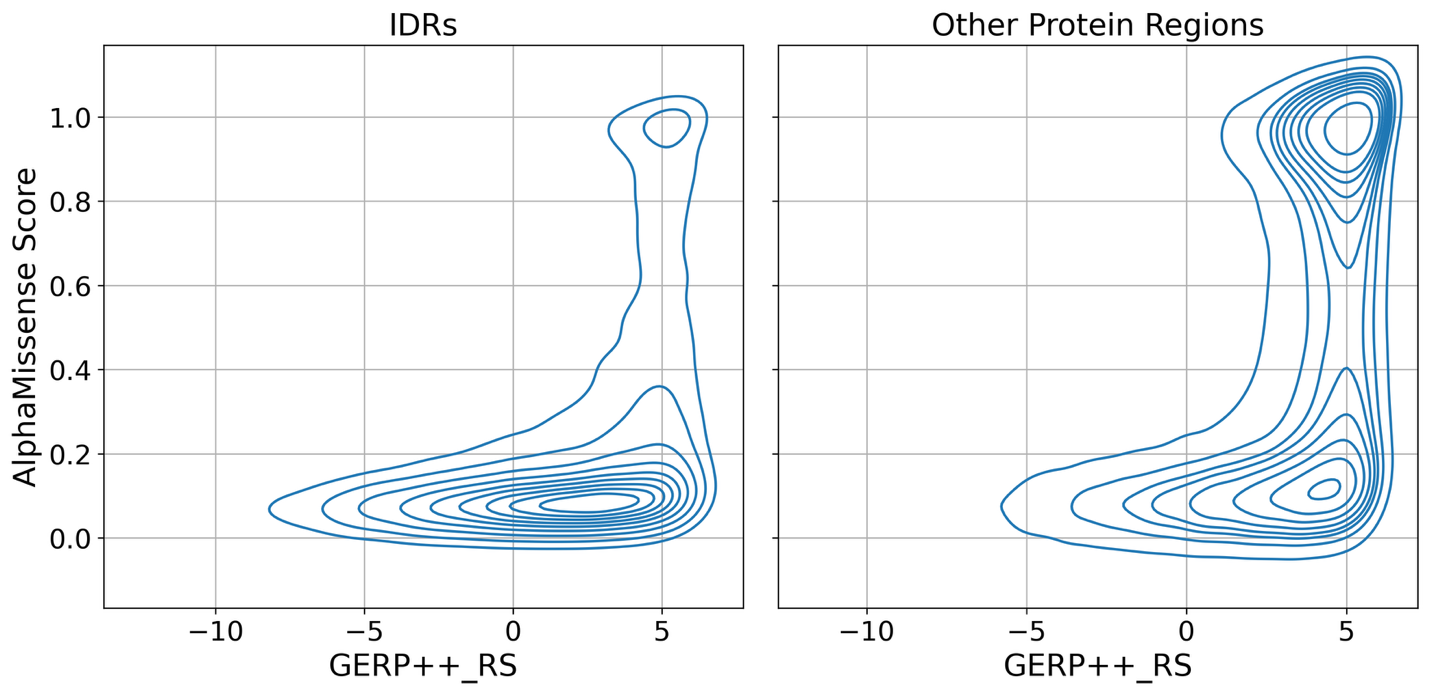

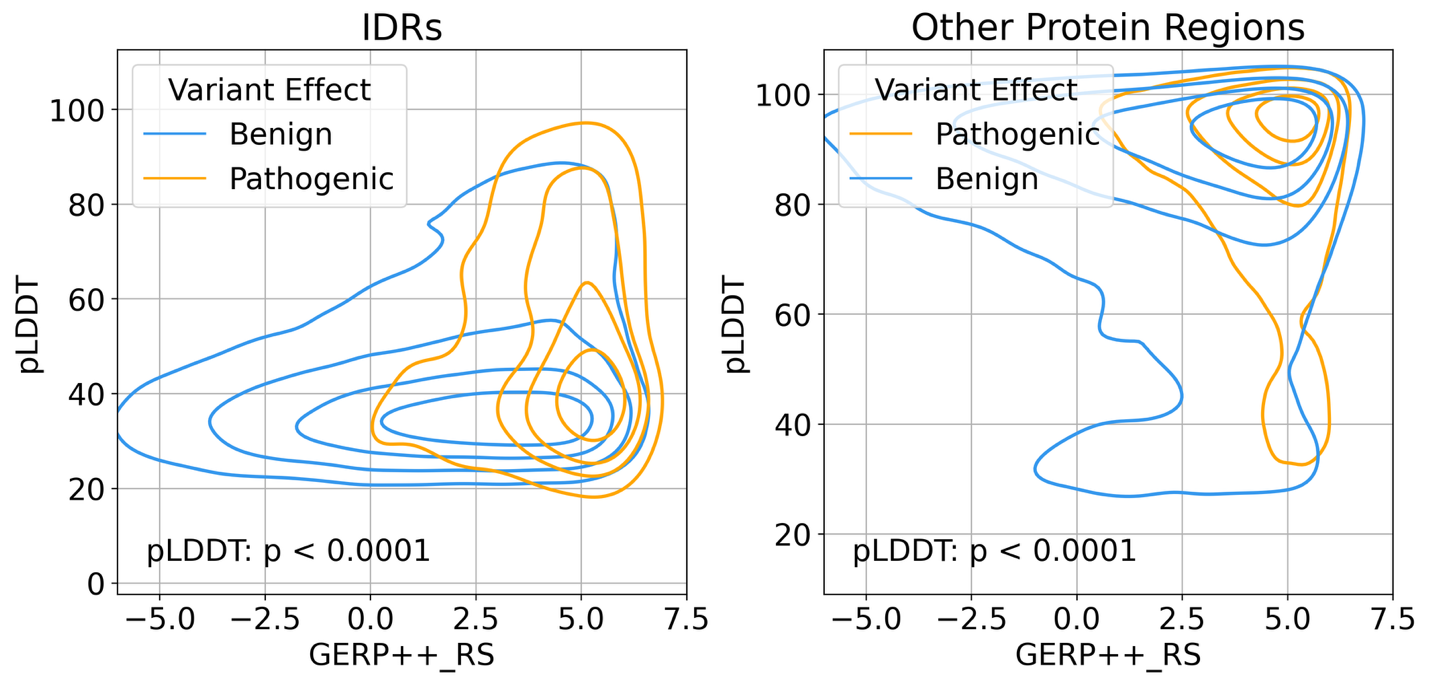

**C**

**Supplementary Figure 1. Intrinsically disordered regions show unique conservation characteristics driving AlphaMissense performance. (A)** IDRs includes Intrinsically disordered regions used in our analysis, and Other Protein Regions includes a set of variants collected from ClinVar, HGMD, PrimateAI, and others for regions that do not fall in our IDR database. IDRs exhibit clearly lower genomic constraint, measured using GERP++ RS score. **(B)** Sites in IDRs that exclusively receive large pathogenicity scores from AlphaMissense have greater genomic constraint. In Other Protein Regions, both pathogenic and benign predictions by AlphaMissense have high genomic conservation. **(C)** Sites in IDRs with pathogenic labels from clinical sources have higher predicted local distance difference test (pLDDT), a proxy of order. For Other Protein Regions, while the difference is significant, there is a qualitatively strong overlap of pathogenic and benign variants in highly ordered regions. Mann Whitney U Tests were used to compute test statistics and p-values.

| **Model** | **Feature Name** | **Position relative to site** | **Mean Decrease in Impurity** |
| --- | --- | --- | --- |
| **MDmis (Res + Pair)** | SASA (Standard Devation) | -3 | 0.017 |
|  | Phi Angle (11^th^ Percentile) | 3 | 0.015 |
|  | Phi Angle (11^th^ Percentile) | -3 | 0.014 |
|  | SASA (Standard Deviation) | 3 | 0.013 |
|  | Phi Angle (10^th^ Percentile) | 3 | 0.013 |
|  | SASA (Mean) | -3 | 0.011 |
|  | Chi Angle (3^rd^ Percentile) | 2 | 0.011 |
|  | SASA (Standard Deviation) | 0 | 0.011 |
|  | Chi Angle (3^rd^ Percentile) | 3 | 0.011 |
|  | Phi Angle (9^th^ Percentile) | -3 | 0.010 |
| **MDmis + MSA + ESM1b** | LLR | 0 | 0.078 |
|  | Entropy of Site | 0 | 0.032 |
|  | Average Entropy of Region | -9 to 9 | 0.023 |
|  | SASA (Standard Deviation) | -3 | 0.013 |
|  | Phi Angle (12^th^ Percentile) | 3 | 0.013 |
|  | Phi Angle (10^th^ Percentile) | -3 | 0.012 |
|  | Phi Angle (11^th^ Percentile) | 3 | 0.0099 |
|  | SASA (Standard Deviation) | 3 | 0.0088 |
|  | SASA (Mean) | -3 | 0.0087 |
|  | Conservation of sequence composition | -9 to 9 | 0.0087 |

**Supplementary Table 1. Feature importances for trained Random Forest models using features derived from molecular dynamics.** Feature importances are computed as mean decrease in impurity and ranked to show the top 10 features across all five training folds. The position is provided relative to site of mutation for residue features. LLR refers to ESM1b’s log-likelihood ratio output that is passed as a feature in our model.

**A**

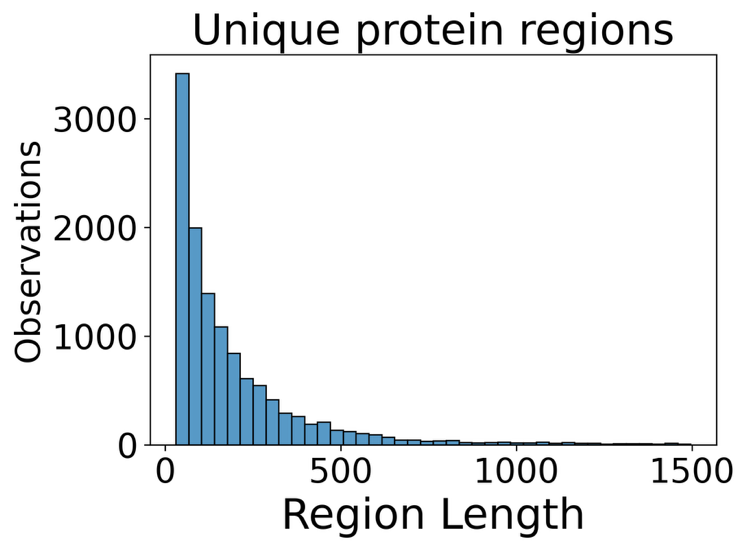

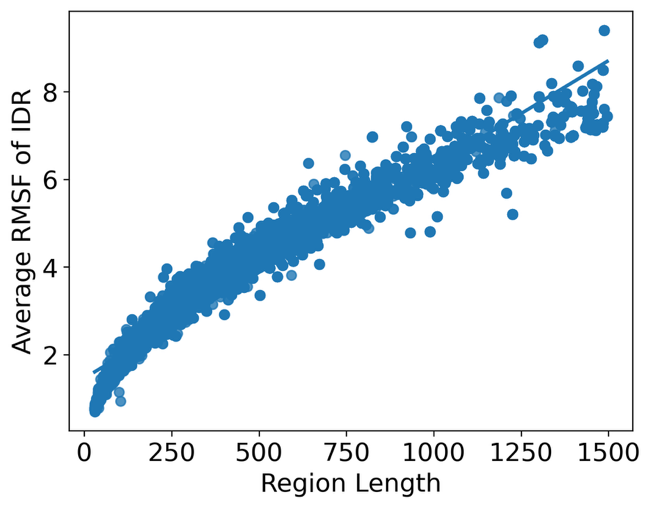

**C**

**B**

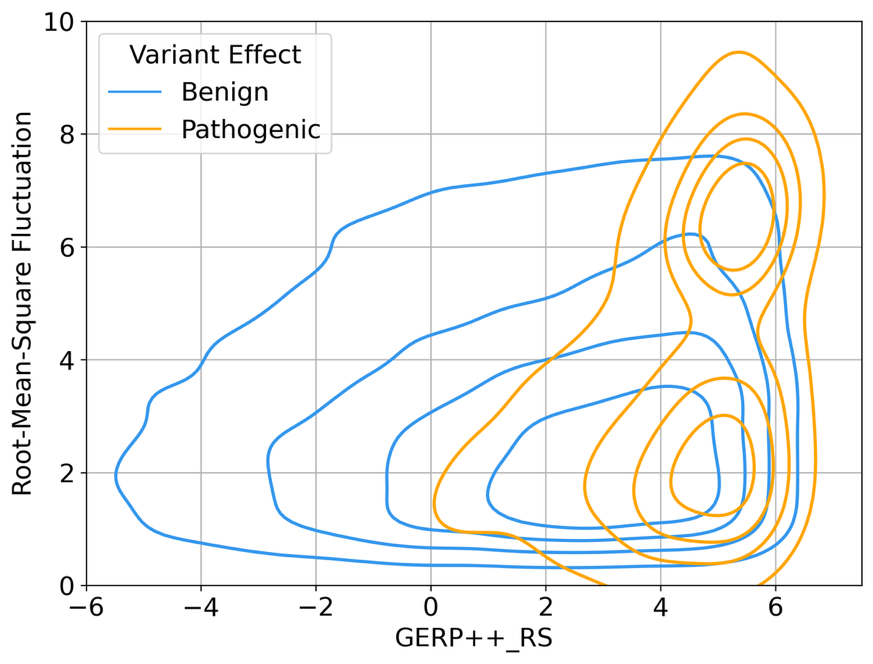

**Supplementary Figure 2. Root-Mean-Square Fluctuation reveals length bimodality in pathogenic variants. (A)** Histogram of length of IDRs in our data without considering variant labels shows a highly right skewed distribution with very few IDRs beyond 500 amino acids. **(B)** Root-mean-square fluctuation (RMSF) measures the fluctuation of residue positions between frames in the simulation. RMSF was averaged for the entire IDR and compared with the length of the IDR. We observe a clear linear relationship between the two, indicating that due to the large spread of IDR length, RMSF is not reflecting fluctuation but rather a technical artifact of length. **(C)** We found that pathogenic variants show a clear bimodal distribution of RMSF, with two peaks separated by a valley at 4.5 Angstroms. This second peak represents a small number of lengthy IDRs with many pathogenic variants.

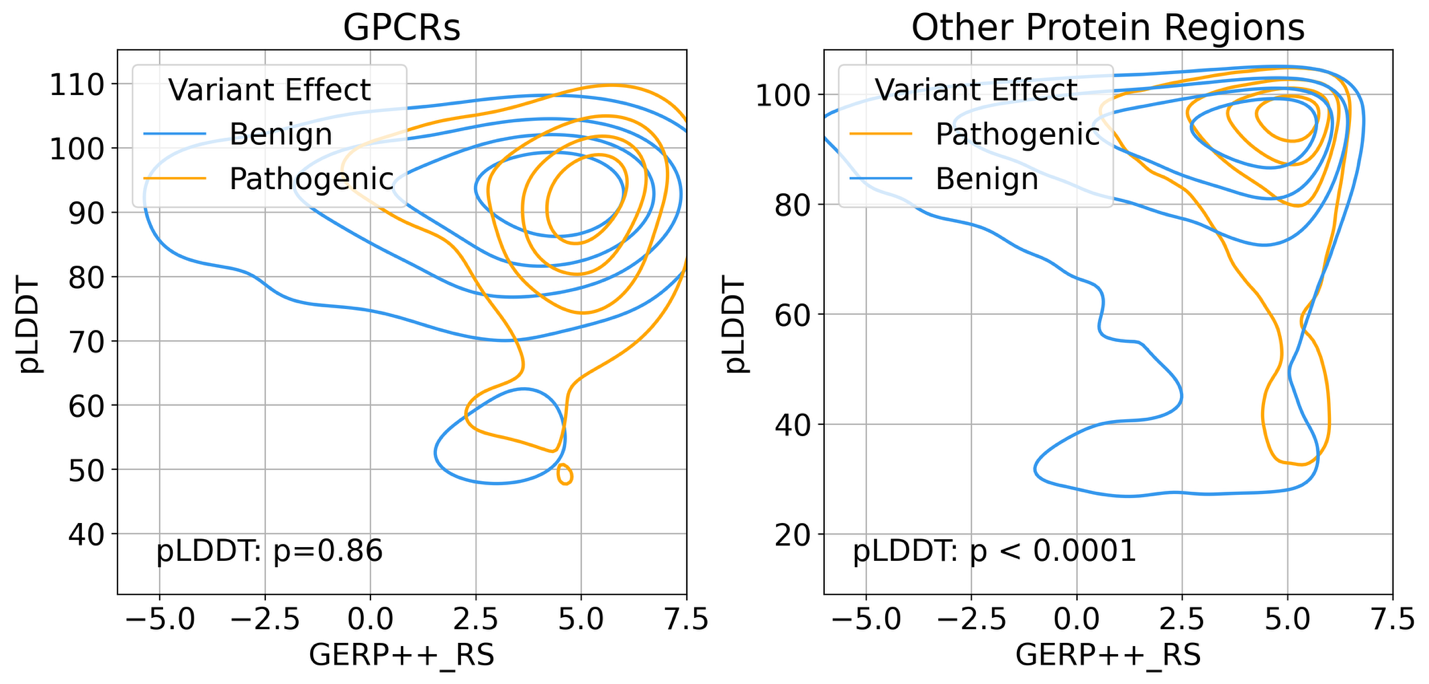

**B**

**C**

**A**

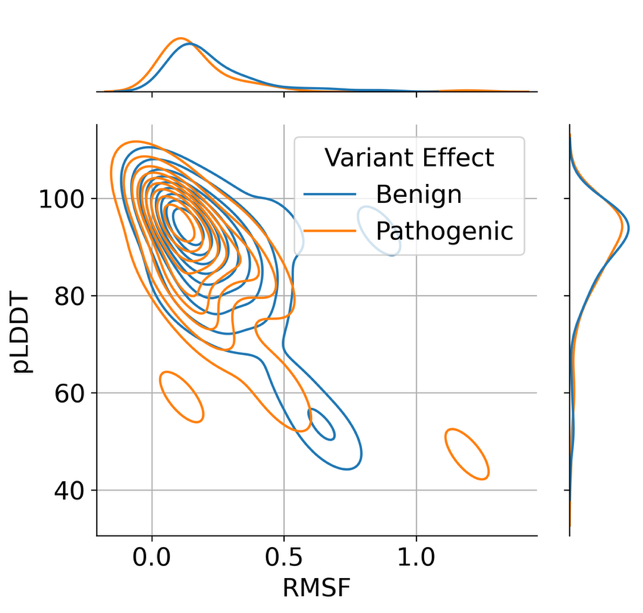

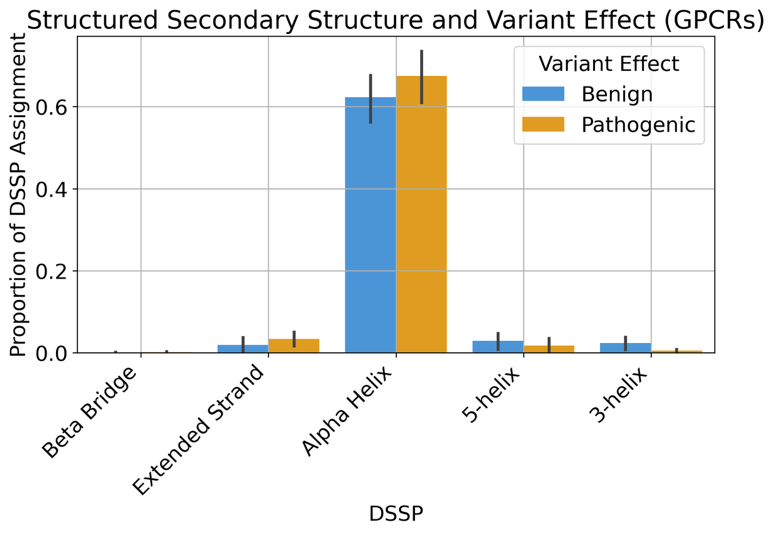

**Supplementary Figure 3. MD simulations of GPCRs reflect different patterns compared to IDRs.** GPCRmd database contains molecular dynamics simulations of GPCRs and contains primarily ordered protein regions. **(A)** Predicted local distance difference test (pLDDT) scores from AlphaFold2 serves as a proxy for order. Benign and pathogenic variants G-protein coupled receptors (GPCRs) have similar pLDDT values. In other protein regions, which contain a mix of primarily ordered domains and some disordered sites, the pLDDT scores show a qualitatively clear overlap between pathogenic and benign variants. **(B)** pLDDT scores plotted with respect to root-mean-square fluctuation of residues in GPCRmd show inverse correlation. **(C)** DSSP secondary structure assignments of residues in GPCRmd are shown with their proportions during the trajectories in GPCRmd database. There are no significant differences in proportions between pathogenic and benign variants. Additionally, all residues with variants appear to spend greater than 60% of their simulations on average as alpha-helices.

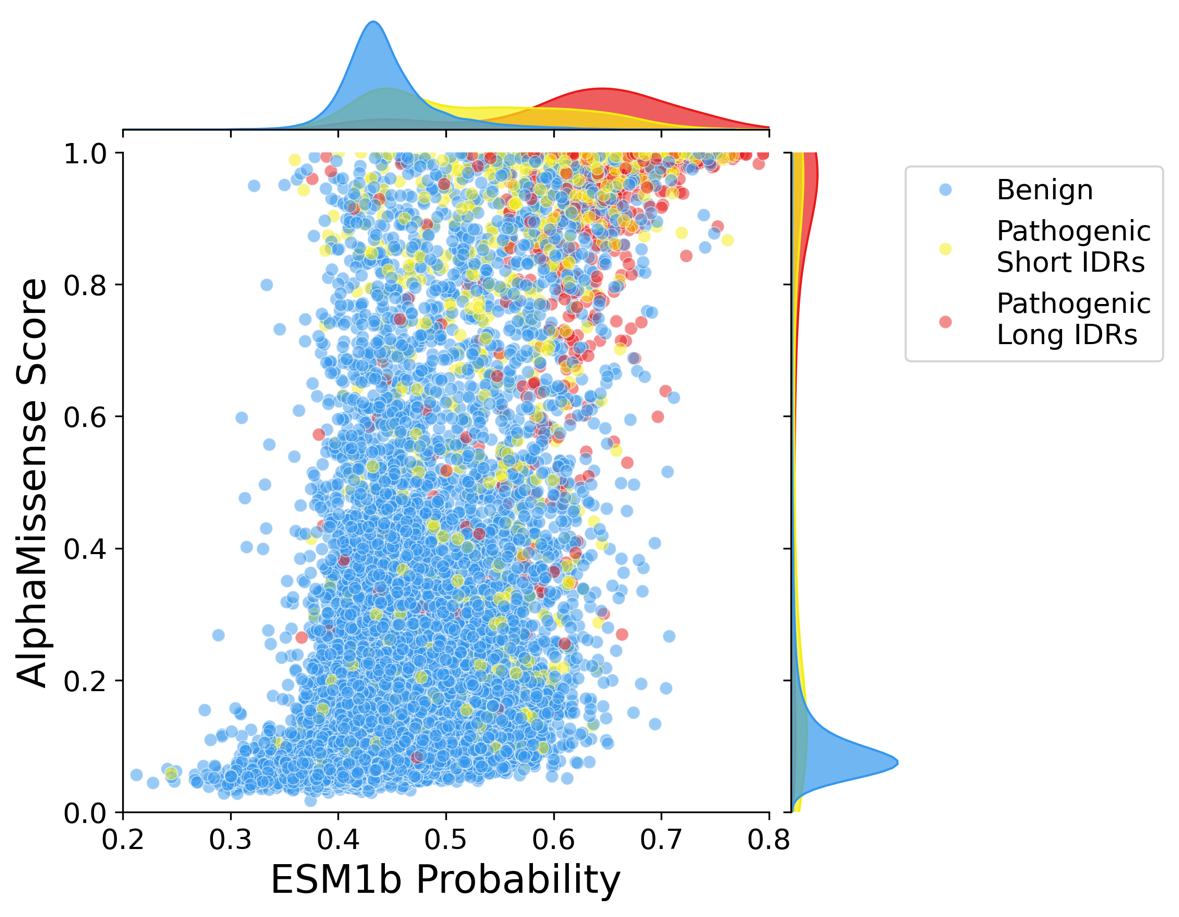

**Supplementary Figure 4. Comparison of ESM1b and AlphaMissense probabilities shows patterns of disagreement.** ESM1b scores and AlphaMissense scores were compared using a scatterplot with their marginal kernel density functions, colored by variant classifications. ESM1b probabilities are obtained by min-max scaling log-likelihood ratio values over the entire human proteome.

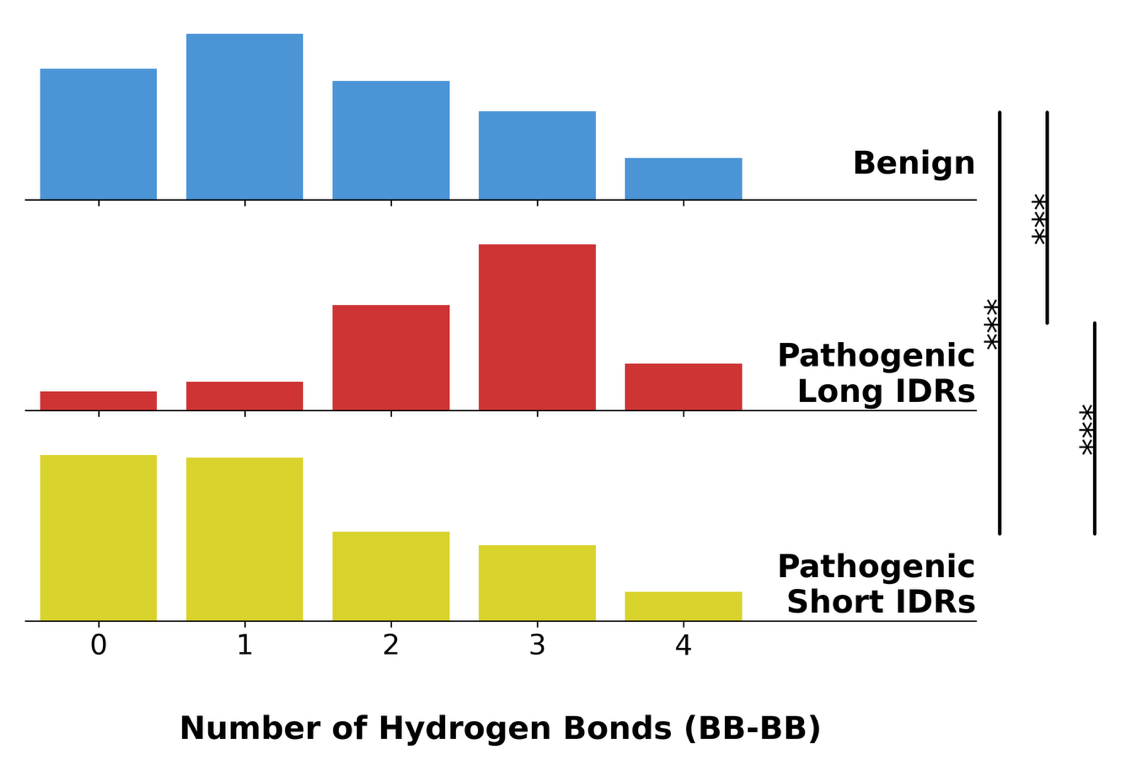

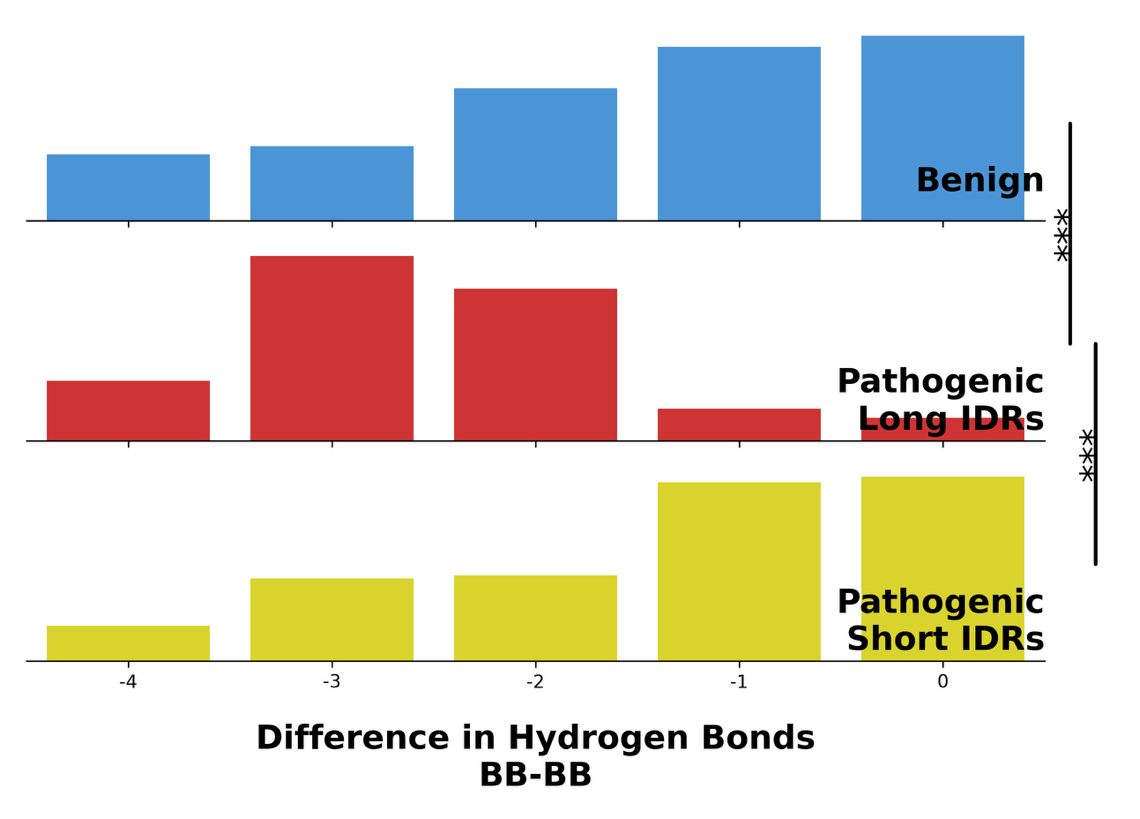

**Supplementary Figure 5. Pathogenic variants in Long IDRs show mode at 3 backbone-backbone hydrogen bonds that disappear after missense variant. (A)** Barplot shows proportion of variants in Long IDRs that have backbone-backbone hydrogen bonds for more than 40% of the trajectory. There is a strong peak at 3 interactions for pathogenic variants in Long IDRs. **(B)** When compared to wild-type simulations, mutated MD simulations show a statistically significant decrease in number of hydrogen bonds between backbones, with loss of 3 interactions having the greatest proportion. Significance stars: p<0.0001: ***, p<0.001: **, p<0.05: *

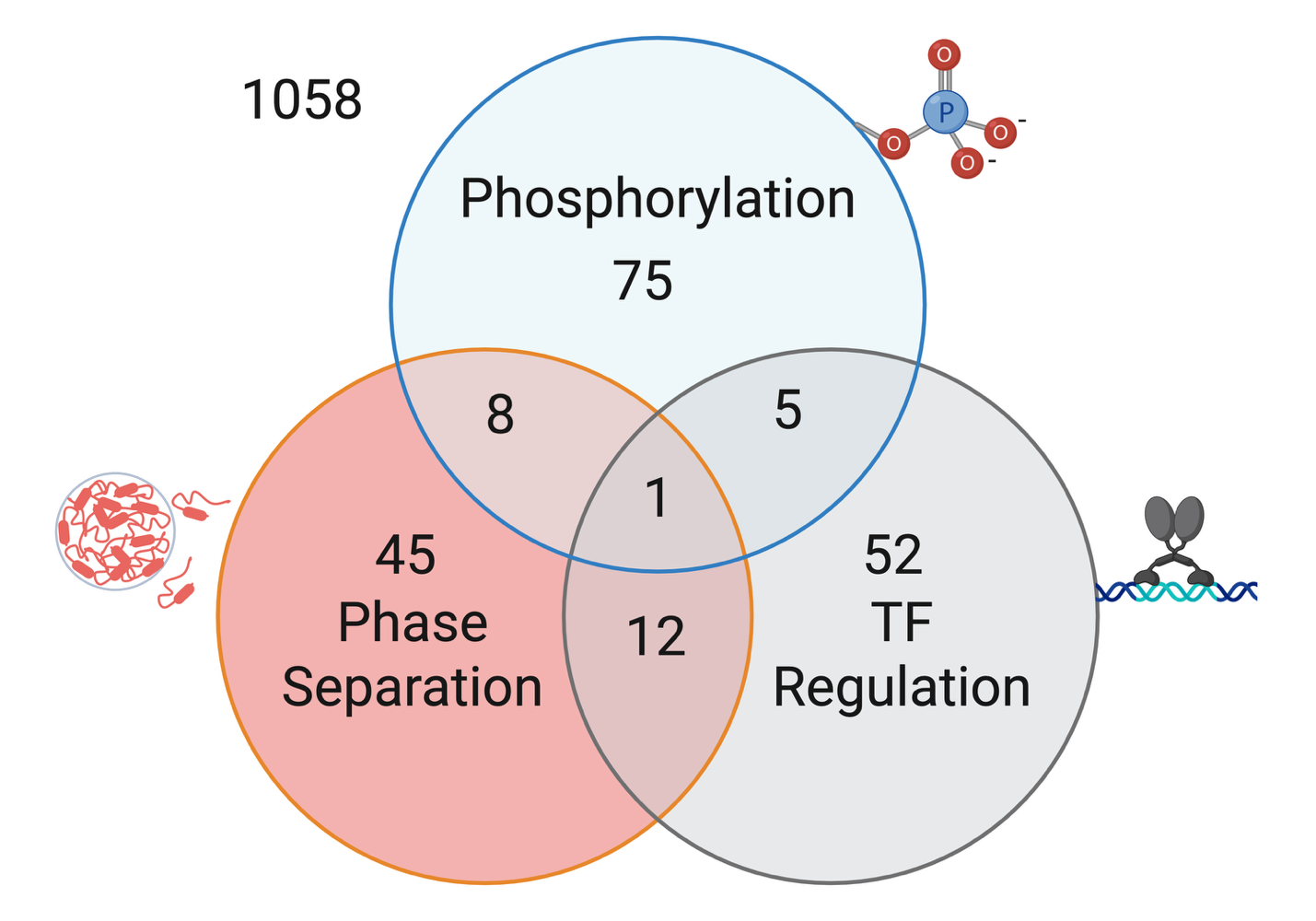

**Supplementary Figure 6. Venn diagram illustrating overlap between functional annotations.** Using dbPTM, TFRegDB, and PhaSepDB, we isolate sites with pathogenic variants in Short IDRs and compare numbers of overlapping variants. The three functional annotations imply little overlap overall. The most sites are shared between transcriptional regulation and phase separation.
